# Differential translocation of bacteriophages across the intestinal barrier in health and Crohn’s disease

**DOI:** 10.1101/2024.09.17.613249

**Authors:** Clara Douadi, Ilias Theodorou, Quentin Lamy-Besnier, Olivier Schiettekatte, Yanis Sbardella, Loïc Brot, Paolo Emidio Costantini, Roberto Saporetti, Alberto Danielli, Matteo Calvaresi, Marianne De Paepe, Harry Sokol, Diego Garcia-Weber, Véronique Carrière, Sophie Thenet, Luisa De Sordi

**Affiliations:** Sorbonne Université, INSERM, Centre de Recherche St Antoine, 75012 Paris, France; Paris Centre for Microbiome Medicine (PaCeMM) FHU, Paris, France; Université Paris-Saclay, INRAE, AgroParisTech, Micalis Institute, Jouy-en-Josas, France; Dipartimento di Farmacia e Biotecnologie (FaBiT) – Alma Mater Studiorum - Università di Bologna, Bologna, Italy; IRCCS Azienda Ospedaliero-Universitaria di Bologna, Bologna, Italy; Dipartimento di Chimica “Giacomo Ciamician” – Alma Mater Studiorum - Università di Bologna, Bologna, Italy; AP-HP, Saint Antoine Hospital, Gastroenterology Department, 75012 Paris, France; EPHE, PSL University, Paris, France

**Keywords:** Intestinal barrier, barrier defect, intestinal permeability, bacteriophages, phages, inflammation, epithelial barrier, endothelial barrier, inflammation, Crohn’s disease

## Abstract

Impaired intestinal barrier function is a major feature of Crohn’s disease (CD), leading to exacerbated inflammation in response to the microbiota. In this context, the translocation of intestinal bacteriophages (phages) and their effects on the host have been little investigated. We used phage fluorescence imaging coupled with ex-vivo and in-vitro models that mimic physiological and inflammatory conditions and found that phages can translocate across the intestinal barrier without disrupting its integrity. Although the translocation rate across the intestinal epithelium depended on phage morphology and the condition of the barrier, these factors did not influence the crossing of phage across the vascular endothelium. Virome analysis confirmed that viral sequences shared between blood and stool samples are more abundant in CD patients than healthy subjects, indicating that a barrier defect facilitates phage translocation from the gut to the bloodstream.

## INTRODUCTION

The intestinal mucosa plays a crucial role in preventing the invasion of luminal microbiota, acting as an essential barrier function for the body, in addition to nutrient absorption and endocrine functions^1^. The maintenance of intestinal barrier integrity results from the combined efforts of specialized epithelial and endothelial cells, as well as immune cells and accessory cells such as enteric glial cells and pericytes. The epithelial barrier consists of a monolayer of tightly sealed differentiated cells lining the gut lumen, that are coated with a layer of mucus and maintained together by apical junctional complex comprising tight junctions, adherens junctions, and desmosomes^2^. The endothelial barrier, forming a deeper layer of protection located in the mucosa and submucosa, is composed of endothelial cells that line the interior of intestinal capillaries and blood vessels^3^. These cells are interconnected by tight and adherens junctions, creating an additional obstacle to be crossed to reach the bloodstream and distant organs^4^. Both the epithelial and endothelial barriers are semi-permeable and essential for regulating the exchange of molecules and fluids from the intestinal lumen into the circulation. An imbalance in their structure can allow unrestrained crossing of molecules and micro-organisms from the intestinal lumen, triggering an excessive immune reaction. An impaired intestinal barrier has been reported in a wide array of human diseases, both extra-intestinal^5–8^, such as multiple sclerosis and non-alcoholic fatty liver disease, and intestinal, including inflammatory bowel disease (IBD).

Crohn’s disease (CD), one of the two main types of IBD, is characterized by chronic relapsing inflammation of the gastro-intestinal tract (GIT), leading to bowel damage and a reduced quality of life for patients^9,10^. CD is considered to be due to an abnormal immune response against the gut microbiota in patients with a genetic predispositions and influenced by environmental factors, resulting in chronic inflammation^11–13^. In addition, CD is associated with a local loss of the mucus layer^14^, increased intestinal permeability^14–17^, and endothelial dysfunction^18,19^. These harmful defects facilitate the passage of bacteria or their products from the GIT into systemic circulation^20–23^. Furthermore, the increased recruitment of leukocytes in the lamina propria may lead to the local accumulation of proinflammatory mediators, such as inflammatory cytokines, further exacerbating intestinal barrier failure^24,25^.

Bacteriophages (phages) are the most abundant viruses of the gut microbiota, representing between 10^9^ to 10^10^ virus-like particles (VLP) per gram of feces^26^, and are strongly suspected to have a role in shaping gut bacterial communities^27^. Although several early metagenomic studies reported that the gut phageome differs between CD patients and healthy subjects (HS)^28–30^, it has more recently been shown that the intestinal phageome lacks reliable markers for CD due to its immense diversity and interpersonal variability^31,32^. Phages have been shown to interact with mammalian cells, including intestinal epithelial cells, macrophages and endothelial cells^33–38^. Nonetheless, their translocation across the gut barrier and their effects on its function and inflammation remain largely unknown.

Here, we investigated the interaction of phages with the intestinal barrier, under both physiological and inflammatory conditions. Using ex-vivo and in-vitro models, we specifically addressed whether phages of varying sizes and structures can translocate from the gut to the bloodstream and whether they, by themselves, induce deleterious effects such as hyperpermeability, inflammation, and cytotoxicity. We demonstrate that a relevant proportion of all tested phages can translocate across intestinal epithelium and endothelium without disrupting the integrity of these barriers or triggering an inflammatory response. In addition, analysis of the viral sequences shared between the blood and stool per participant indicates higher phage translocation from the gut to the blood in CD patients.

## MATERIALS AND METHODS

### Patients

Patients diagnosed with CD and healthy subjects were enlisted from the Gastroenterology Department at Saint-Antoine Hospital in Paris, France. The study received approval from the competent ethics committee (Comité de Protection des Personnes Ile-de-France IV, IRB 00003835 Suivithèque study; registration number 2012/05NICB), and all participants provided their consent after being fully informed. Our cohort consisted of 15 CD patients and 14 healthy controls. None of the participants had received antibiotic treatment within the three months preceding sample collection or undergone intestinal surgery. The median age of the cohort was 38 ± 11 years. Further details of the cohort are presented in **Table S1**.

### Sample collection

Blood was drawn into plasma collection tubes containing the anticoagulant EDTA (Dutscher) and promptly placed on ice, followed by centrifugation at 2,000 x g for 10 min to isolate the plasma, which was then filtered through a 0.45-µm cellulose acetate filter (VWR Avantor) and stored at −20°C. Stool samples were gathered directly into RNAlater (Sigma-Aldrich), which precipitates phage particles, and homogenized by vortexing with glass beads in a low-binding Eppendorf tube (Sigma-Aldrich). Following centrifugation at 20,000 x g for 15 min at 4°C, the supernatant was discarded, and the pellet was directly stored at −80°C. Both blood and stool samples from each participant were collected on the same day. For negative controls, sterile water was placed in a blood collection tube and in a low-binding tube.

### Bacterial strains and phage preparation

The bacterial strains used in this study, *Escherichia coli* K-12 MG1655 F+^39^, *Escherichia coli* K-12 MG1655^40^, and *Escherichia coli* C+ (ATCC 8739) were cultured in Luria-Bertani (LB) media (10 g/L tryptone, 5 g/L yeast extract, 5 g/L NaCl) at 37 °C with shaking overnight and used to amplify M13yfp^39^, T4 and ΦX174 phages, respectively, which were titered by plaque assay (**Table 1**). Briefly, infections were initiated by adding phages to bacteria in the early exponential phase (OD_600nm_=0.2) at a multiplicity of infection (MOI) of 0.01. To enhance phage adsorption and amplification, LB media was supplemented with 1 mM CaCl_2_ and MgCl_2_. After overnight incubation and lysis of the bacteria by the phages, remaining bacteria and debris were removed by centrifugation for 30 min at 3,000 x g, and the supernatant filtered sequentially through 0.45-µm and 0.2-µm cellulose acetate filters (VWR Avantor). Phages were concentrated by tangential flow filtration using a Vivaflow® cassette system (Sartorius Stedim Biotech) and stored in a final solution of salt-magnesium (SM) buffer (17 mM MgSO_4_·7H_2_O, 100 mM NaCl, 5 mM Tris-HCl (pH 7.5)) at 4°C.

**Table 1.** Characteristics of the phages used in the study.

| Phage | Family | Size | Bacterial host |
| --- | --- | --- | --- |
| **T4** | *Straboviridae* | Length : 200 nm<br>Width : 90 nm | *E. coli* K-12 MG1655 |
| **M13** | *Inoviridae* | Length : ± 900 nm<br>Width : 7 nm | *E. coli* K-12 MG1655 F+ |
| **ΦX174** | *Microviridae* | Diameter : 25 nm | *E. coli* C+ |

Lysates from the same bacteria cultured without phages were used as a mock negative control and were generated through a series of freeze/thaw cycles followed by sonication to closely mimic phage-induced lysis. The bacterial lysate was subsequently plated on LB agar plates to confirm the death of the bacteria. This sample then underwent the same processing steps used for phage purification.

### Endotoxin removal and determination of the concentration

To effectively remove lipopolysaccharide (LPS), each preparation of purified phage or mock negative control were purified twice using Pierce^TM^ high-capacity endotoxin removal resin (Thermo Fisher Scientific) and then three times using an EndoTrap® HD kit (Lionex) according to the manufacturer’s instructions. The final LPS concentration was determined using a Pierce^TM^ Lyophilized Amebocyte Lysate (LAL) chromogenic endotoxin quantitation kit (Thermo Fisher Scientific) and was shown to be below 1 EU/mL when added to mammalian cells.

### Ex-vivo assays with Ussing chambers

Animal experiments were approved by the ethics committee (Comité National de Réflexion Ethique sur l’Expérimentation Animale) under project n. Ce5/2020/004. Ten- to 12-week-old C57BL/6J male mice were obtained from and housed at the Saint-Antoine Research Center (Plateforme d’Hébergement et d’Expérimentation Animale; Centre De Recherche Saint-Antoine; INSERM UMR_S938) and fed ad libitum until sacrifice.

Intestinal sections (terminal ileum and distal colon) were longitudinally excised and mounted in Ussing chambers (WorldPrecision Instruments, UK) as previously described^40^. Both the apical and basal chambers were filled with 2 ml Ringer buffer solution (115 mM NaCl, 25 mM NaHCO_3_, 1.2 mM MgCl_2_, 1.2 mM CaCl_2_, 2.4 mM K_2_HPO_4_, 0.4 mM KH_2_PO_4_) and maintained at 37°C in a 95% oxygen/5% CO_2_ atmosphere. Following a 20 min equilibration period, phages and FITC-Dextran 4 kDa (FD4) (final concentration: 2 mg/mL; TdB Consultancy) were introduced into the apical chamber. Intestinal permeability was assessed by measuring the fluorescence of FD4 in the basal compartment every 15 min, using a microplate fluorometer (BMG Labtech®). Permeability values are expressed at each time point as the percentage of the amount of FD4 initially introduced into the apical chamber. To evaluate phage translocation, the apical chamber was inoculated with purified phages at the beginning of the experiment and phages from the basal compartment counted by plaque assay after 90 min, as detailed in the phage translocation assays in the “cell models” section.

### Cell culture and treatments

The Caco-2/TC7 cell line is a clonal population of human colon carcinoma-derived Caco-2 cells^42,43^. Caco-2/TC7 cells were cultured on six-well Transwell® 3 μm-pore size filters (Sarstedt) for 19 days to obtain fully differentiated enterocyte-like cells as previously described^44^. Briefly, cells were cultured in high-glucose Dulbecco’s modified Eagle’s medium Glutamax (Thermo Fisher Scientific) supplemented with 20% heat-inactivated fetal calf serum (FCS) (Sigma-Aldrich), 1% non-essential amino acids, penicillin (100 IU/ml), and streptomycin (100 μg/ml) for 7 days and then switched to asymmetric conditions, i.e., with medium containing 20% FCS in the basal compartment and serum-free medium in the apical compartment. Human umbilical vein endothelial cells (HUVECs) were obtained directly from umbilical cords donated by Bluets Maternity Hospital (Pierre Rouquès Hospital, Paris - Dr. Jessica Dahan Saal) or purchased from Lonza. These cells were cultured in flasks pre-coated with 10 µg/ml human fibronectin (Sigma-Aldrich) in endothelial basal medium (EBM-2) (Lonza), enriched with 2% fetal bovine serum (FBS) and endothelial cell growth supplement EGM-2 (BulletKit^TM^, Lonza). Before the addition of phage and/or treatments, confluent HUVECs were starved for 2 h in EBM-2 medium supplemented with 1% fetal calf serum without growth supplements (starvation medium) to stop cell proliferation. The cells were seeded in 24-well plates or on 12-well Transwell® 3 μm-pore size filters (Sarstedt) and used up to passage 4.

To increase paracellular permeability, Caco-2/TC7 cells were treated with 5 mM EGTA (Sigma-Aldrich) in the apical compartment or human interferon-γ (IFNγ) and tumor necrosis factor-α (TNFα) (50 ng/mL each; Bio-Techne) in the basal compartment. The EGTA treatment was applied 2 h before adding the phages, whereas TNFα and IFNγ were applied 24 h before. HUVECs were treated with 10 ng/mL TNFα and IFNγ (Bio-Techne) added to the apical compartment of Transwell® filters 18 h before phage addition.

### Phage translocation assays in cell models

Phage translocation was evaluated by inoculating the apical side of the insert with between 10^7^ and 10^9^ PFU of purified phages at the beginning of the experiment. At each timepoint, 100 µL of medium from the basal side was collected to count the translocated phages by plaque assay. To count phages associated with the cells, Caco-2/TC7 or HUVECs were washed twice with phosphate buffered saline (PBS) to remove free phages and the cells were lysed by incubation in 1% Triton X-100 (Sigma-Aldrich) for 5 min and 100 µL of cell lysate was collected.

Samples of basal medium and cell lysates were serially diluted in SM buffer, plated on LB agar plates overlaid with phage-specific *E. coli* strains (see above and **Table 1**), and the plates incubated at 37°C overnight to determine the number of plaque-forming units (PFU).

A mock negative control containing purified lysed bacteria without phages was also added to the cells to confirm that the observed effects were solely due to the presence of the phages (data not shown). In all experiments, the negative control consisted of cell culture medium only.

### Permeability measurements

The paracellular permeability of Caco-2/TC7 and HUVECs monolayers was assessed by introducing 1 mg/mL 4 kDa FITC-dextran (TdB Consultancy) into the apical medium. Samples from the basal medium were collected at various timepoints, and the fluorescence measured at an excitation wavelength of 485 nm and an emission wavelength of 550 nm using a FLUOstar Omega microplate fluorometer (BMG Labtech), as previously described^45^. Trans-epithelial/trans-endothelial electrical resistance (TEER), which inversely correlates with ion permeability, was evaluated both before and after the addition of phages and/or treatment using a Volt-Ohm Meter (Millipore).

For HUVECs, permeability was also monitored in real time using the RTCA-DP xCELLigence system (Agilent), housed in a humidified incubator at 37°C with 5% CO2. The instrument measures electrical impedance on microelectrode-coated plates (Agilent) to track changes in barrier function. It was set to a single frequency of 10 kHz with 3-min measuring intervals over the course of 48 h.

### Cytotoxicity assay and IL-8 secretion

At the end of the experiments, cytotoxicity was assessed by measuring lactate dehydrogenase activity in the apical medium, following the manufacturer’s instructions (Cytotoxicity Detection KitPLUS, Sigma-Aldrich). In addition, the level of the chemokine IL-8 in the basal medium of Caco-2/TC7 cells was quantified using an enzyme-linked immunosorbent assay (ELISA) kit (R&D Systems).

### Preparation, purification and conjugation of CF488 fluorophore to M13 phages

M13 phage was produced by infecting *E. coli* TG1 grown to an OD_600nm_=0.4 with 10^11^ PFU of M13KO7 helper phages (New England BioLabs) for 2 h. After infection, 10 mL was transferred to a flask containing 400 mL LB supplemented with kanamycin (25 mg/L) and the flask was incubated for 24 h at 37°C with shaking. Phage purification was performed as previously described^46^. The bacterial culture was centrifuged at 12,000 x *g* for 30 min. The supernatant was supplemented with PEG 8000 (4% w/v) and NaCl (3% w/v) and incubated at 4°C for 90 min. The solution was then centrifuged for 15 min at 12,000 x *g* and the phage pellet was resuspended in PBS before a further purification step by isoelectric point (IEP) precipitation^47^. Purified phages were finally resuspended in PBS. Phage concentrations were determined by measurement of the absorbance at a wavelength of 269 nm in a UV-Vis spectrophotometer (extinction coefficient ε = 3.84 cm^2^/mg) using a 320 nm wavelength spectrum as baseline.

An aliquot of 1 µmol CF™ 488A succinimidyl ester (CF488-SE; Biotium) was dissolved in 100 µL anhydrous DMSO to obtain a 10 mM CF488-SE solution. Then, 29.3 µL of the CF488-SE solution was added dropwise, with stirring, to 1 mL of 40 nM M13 phages in 100 mM sodium carbonate-bicarbonate buffer (pH 9). The reaction was carried out for 3 h at 25°C with shaking (ThermoMixer HC, S8012-0000; STARLAB, Hamburg, Germany). The CF488-tagged M13 phages were extensively dialyzed against 100 mM sodium carbonate-bicarbonate buffer (pH 9) using a regenerated cellulose membrane (14 kDa cut-off) to remove reaction by-products. The last dialysis cycle was performed against 10 mM PBS (pH 7.4).

### Visualization of the endocytic pathway in HUVECs

To evaluate the endocytic process, transferrin-Cy3 (3.5 µg/mL; Cy3 ChromPure Human transferrin; Jackson ImmunoResearch Europe) was added to HUVECs simultaneously with the phages. Acidic compartments were visualized by staining the cells with Lysotracker® Deep Red (200 nM; Thermo Fischer Scientific) either 30 min or 1 h prior to fixation.

### Confocal microscopy

Caco-2/TC7 cells were fixed with 4% paraformaldehyde for 30 min and then permeabilized for 1 h with PBS containing 1% bovine serum albumin (BSA), and 0.5% Triton X-100 (Sigma-Aldrich), and finally blocked with PBS containing 3% BSA. HUVECs were fixed with 4% paraformaldehyde for 15 min, followed by permeabilization for 10 min in PBS containing 2% BSA and 0.1% Triton X-100 (Sigma-Aldrich), followed by blocking with 2% BSA. Cells were then incubated at 37°C with primary antibodies: tricellulin (1:200; MARVELD2 700191; Thermo Fischer Scientific), ZO-1 (1:200; ZO1-1A12; 33-9100 Thermo Fischer Scientific), and E-cadherin (1:500; ECCD2 M108; Takara Bio Europe). Alexa Fluor^TM^ 647, and Alexa Fluor^TM^ 546–conjugated anti–immunoglobulin G were used as secondary antibodies (1:400; Invitrogen, Thermo Fischer Scientific). Nuclei were stained with 4′-6-diamidino-2-phenylindole (DAPI) (1 µg/mL, Invitrogen), and actin with Alexa Fluor^TM^ 647/546-labelled phalloidin (1:400; Invitrogen). Specimens were mounted in fluorescence mounting medium (ProLong^TM^, Gold Antifade, Thermo Fisher Scientific). Confocal laser scanning microscopy was carried out with an Olympus Fluoview FV3000 instrument using a ×60 oil immersion objective. To obtain Z stacks, optical sections were taken over 0.43 μm. Images were acquired by FV3000 software (Olympus) and processed using ImageJ Fiji (http://imagej.net/ij/index.html). M13 phage-positive Caco-2/TC7 cells were analyzed using Qupath. Cells were segmented by setting a threshold of 10 to detect nuclei, followed by an expansion adjustment of 10 µm to define the cell boundaries. The Green Max signal was selected to identify M13 phage particles. Three fluorescence intensity thresholds were then applied to classify cells: 48 for low-positive (+), 100 for medium-positive (+), and 200 for high-positive (+). Cells were categorized based on these thresholds by analyzing 10 randomly selected images.

### Quantification of T4 phages by qPCR

Viral DNA isolation was carried out using a phenol/chloroform protocol adapted from Billaud et al.^48^, and the DNA concentration measured using a Qubit^TM^ 4 Fluorometer (Invitrogen). For qPCR, a 20-µL reaction mix was assembled with 50nM T4 phage primers (forward : 5’-ACTGGCCAGGTATTCGCA-3’, and reverse : 5’-ATGCTTCTTTAGCAGCGGCA-3’), SYBR^TM^ Green PCR Master Mix (Applied Biossystems), and the sample. The qPCR reactions were run on a StepOnePlus^TM^ Real-Time PCR System (Thermo Fisher Scientific) with the following cycling conditions : 95°C for 10 min, 40 cycles of 95°C for 15 s, and 60°C for 1 min, followed by a melt curve setting of one cycle of 95°C for 15 s, 60°C for 1 min and 95°C for 15 s. A calibration curve using T4 DNA (ranging from 3.1 to 1.10^-8^ ng) was used to determine the genome copy/mL of T4 phages in each sample. The conversion of qPCR results to copy number/mL followed the principles outlined by Peng et al^49^.

### Viral DNA sequencing

Viral DNA from both blood and stool samples was isolated using a protocol dedicated to virome extraction adapted from Billaud et al^48^. Both stool samples (0.2 g) and plasma samples (3 mL) were thawed on ice. Stool samples were subsequently suspended in 7 mL 10 mM Tris pH 7.5. All samples were then centrifuged at 4,750 x *g* for 10 min at 4°C, and the supernatant was filtered through a 0.45-µm PES membrane filter to eliminate bacteria. The filtered supernatant was then treated with benzonase-nuclease (250 U) for 2 h at 37°C. Viruses were concentrated by mixing the samples with 0.5 M NaCl and 10% weight/volume PEG-8000, followed by overnight incubation at 4°C. Then, samples were centrifuged at 4,750 x g for 30 min at 4°C, and the resulting precipitate was collected and re-suspended in 400 µL 10 mM Tris pH 7.5. An equal volume of chloroform was added, and the samples were gently shaken to eliminate remaining bacteria and decrease extracellular vesicle contamination. Samples were centrifuged at 16,000 x g for 5 min at 4°C to extract the aqueous phase. To remove non-encapsidated nucleic acids, the samples were treated with 8 U TURBO DNAse I (Ambion/Thermo Fisher Scientific) and 20 U RNAse (Thermo Fisher Scientific). The DNAse was then inactivated with EDTA at a final concentration of 20 mM. To disrupt the capsids, 40 µg proteinase K and 20 µL 10% SDS were added before incubation at 56°C for 3 h. Subsequently, the samples were transferred into tubes containing an equal volume of phenol-chloroform-isoamyl alcohol (25:24:1, Thermo Fisher Scientific). After vigorous vortexing, they were centrifuged at 16,000 x g for 5 min at 4°C. The resulting aqueous phase was extracted and transferred to a new tube. Chloroform/isoamyl alcohol (24:1, Thermo Fisher Scientific) was added, followed by another identical centrifugation to eliminate traces of phenol. The resulting aqueous phase was then extracted, and the DNA was precipitated using 0.3 M sodium acetate, 2.5 volumes of ethanol, and glycogen (20 µg/mL final concentration), followed by a 1-h incubation at −20°C. Samples were then centrifuged at 16,000 x g for 30 min at 4°C, and the pellet was washed twice with 0.5 mL cold 75% ethanol. After ethanol removal, the dry pellet was resuspended in 10 mM Tris pH 8 and stored at −20°C. Libraries were prepared using the xGen kit, which includes a step to convert ssDNA to dsDNA with minimal bias^49^. The resulting libraries were then sequenced on an Illumina apparatus (paired-ended 150 bp reads), generating 9.2 million (± 6 million) pairs of reads (**Table S2**).

### Determination of vOTUs

Sequencing reads underwent cleaning using fastp 0.23.1^50^ with the following parameters: -- trim_front1 2 --trim_tail1 2 --trim_front2 15 --trim_tail2 2 -r -W 4 -M 20 -u 30 -e 20 −l 100. Human reads were eliminated by aligning the cleaned reads against the human genome (GRCh38) using bowtie2 2.4.1^51^ with the --very-sensitive option. Four samples (H7_F, H10_F, H13_B, and CD7_B) were excluded from further analysis due to having less than 100,000 pairs of reads remaining.

Assembly was conducted using the cleaned, non-human reads both per participant (utilizing both blood and stool samples) and collectively from samples of all participants. In both cases, paired-end and unpaired reads were used and deduplicated before assembly using the dedupe function of bbmap 38.86^52^. Assembly was performed using SPAdes 3.14.0^53^ with default parameters, excluding contigs smaller than 2 kb, corresponding to the minimal size of DNA viral genomes.

Contigs were clustered to define operational taxonomic units (OTUs) using an approach adapted from Shah et al^54^. Initially, pairwise alignment of all contigs was performed using BLAT 36^55^. Contigs exhibiting a total alignment length against themselves > 110% were identified as chimeras and removed. Then, clustering was performed at 95% average nucleotide identity (ANI), representing species boundaries, using scripts described in Shah et al^54^.

Two strategies were combined to detect viral OTUs: dedicated tools and sequence homology in virus databases. For the dedicated tools, VirSorter2 2.2.3^56^ was used with specific parameters to enhance stringency: *--include-groups dsDNAphage,NCLDV,RNA,ssDNA,lavidaviridae --exclude-lt2gene --min-length 2000 --viral-gene-required --hallmark-required --min-score 0.9 --high-confidence-only*. VIBRANT 1.2.1^57^ was also used. In addition, CheckV 0.8.1^58^ was applied to all OTUs, with only those surpassing a "medium" viral quality being retained. For the databases, the OTUs were compared to sequences in various virus databases (GPD^59^, GVD^60^, COPSAC^54^, MGV^61^, CHVD^62^, and RefSeq Virus 07/2023^63^) using BLAT 36^55^, considering OTUs with > 90% ANI homology to at least one database as being viral. Finally, the vOTUs from both methods were merged to create a non-redundant list of 17,454 viral vOTUs. Complete information for each vOTU can be found in Lamy-Besnier et al^32^.

### Characterization of the vOTUs

For classification, vOTUs were combined with the INPHARED database^64^ (02/07/2023) before clustering with vContact2 0.9.19^65^ using default parameters. A classification was assigned from the obtained network using graphanalyzer 1.5.1^66^.

Host prediction was conducted using iPHoP 1.2.067 with the associated database (09/2021) using default parameters.

### Ecological Analysis

Cleaned, non-human reads underwent mapping using bwa-mem2 2.2.1^68^ against all OTUs with default parameters. The resulting SAM file was converted to BAM and sorted by read name using samtools^69^ 1.9 before filtering out low-quality alignments using msamtools filter 1.1.3. Alignments were retained if they were > 80 bp long, had > 95% overall identity, and > 80% of the read aligned. Only the best alignment (or alignments in cases of ties) for each read was retained.

Alignments were then converted into counts using msamtools profile, with the multi=proportional option to distribute reads mapping to different OTUs according to their relative abundance. The coverage and depth of each OTU for each sample were determined using msamtools coverage. OTUs with coverage < 50% of the OTU size or an average depth < 1 in a given sample were removed.

For viral community analysis, vOTU counts were extracted. vOTUs present in negative control samples were removed from the other samples. Counts were then normalized by vOTU length and transformed into relative abundance for comparison across vOTUs and samples, forming the primary abundance table for analysis. This relative abundance table was analyzed using the R package phyloseq^70^ 1.46.0.

### Statistical analysis

Statistical analyses were performed using GraphPad Prism 6.0 (GraphPad Software). The assumption of normality was assessed using the Shapiro–Wilk test. Two group comparisons were performed using Student’s t-test (parametric) or Mann-Whitney test (nonparametric) and comparisons involving multiple groups were carried out using one-way analysis of variance (ANOVA) or Kruskal–Wallis test when the assumptions of ANOVA were not met. Spearman’s test was used to assess the correlation between the two variables tested. A level of p < 0.05 was considered significant.

## RESULTS

### Phages cross the entire mouse intestinal wall

We investigated the ability of phages to cross entire intestinal tissues by initiating experiments using the Ussing chamber system. The translocation capacity of T4, M13 and ΦX174 phages, each belonging to distinct families with different structures and sizes (**Table 1**), was measured in the basal (serosal) compartment 90 min after their addition to the apical (luminal) compartment, along with the fluorescent tracer FITC-Dextran 4 kDa (FD4) (**Figure 1A**). The rate of FD4 passage averaged 0.03% for the ileum and 0.018% for the colon, indicating that both intestinal tissues remained viable and undamaged throughout the experiment (**Figure 1B**). As expected^72–74^, the ileum was approximately twice as permeable as the colon (**Figure 1B**). Remarkably, 10^2^ to 10^4^ PFU of all tested phages demonstrated the capacity to cross the mouse ileum sections, retaining the ability to infect bacteria and form plaques (**Figure 1C-E**), with translocation rates ranging from 0.000093% to 0.00041% (**Table S3**). Translocation of functional T4 and M13 phages was also observed across the colon (**Figure 1C-D**), but not that of ΦX174 (**Figure 1E**). These data provide evidence that phages can functionally translocate across the entire intestinal tissue.

**Figure 1:**
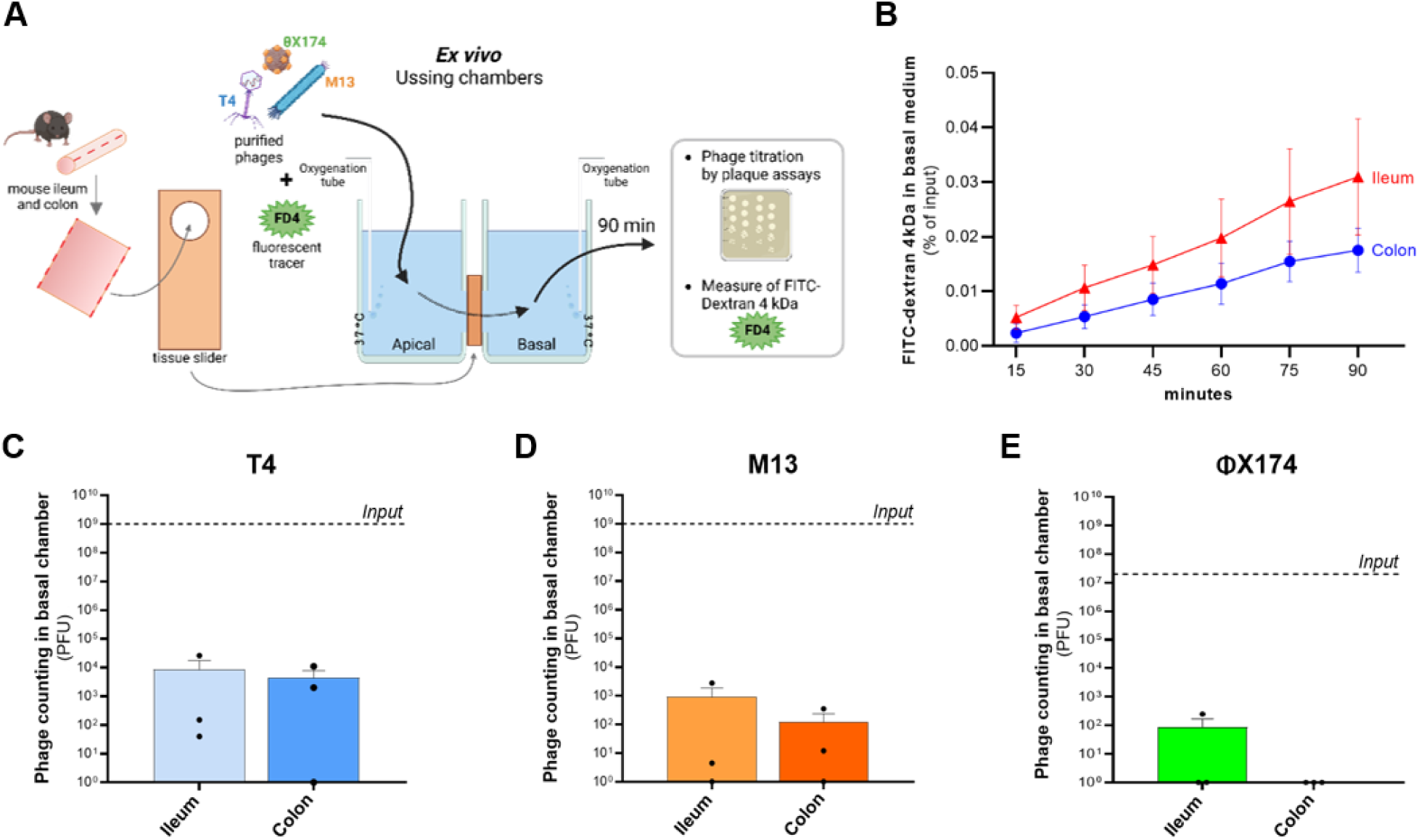
Translocation of phages across intestinal wall in healthy mice. **(A)** Overview of Ussing chamber experimental setup. **(B)** Level of FITC-Dextran 4kDa in basal medium over 90 min for ileum (red curve) and colon (blue curve) tissues from healthy mice. Data are presented as the percentage of fluorescent tracer added in the apical chamber over time. For each intestinal segment, six replicates were performed. **(C-E)** Translocation of **(C)** T4, **(D)** M13 or **(E)** ΦX174 phages across mouse ileum and colon tissues after 90 min of phage addition in the apical compartment. For each phage, three ileum or colon tissue samples were taken from two or three individual mice. Data are represented as mean ± SEM.

### Phages translocate across intestinal epithelial cells without disrupting barrier integrity

As the epithelial barrier is the first to be encountered by the intestinal microbiota, we studied the interaction of phages with this barrier using an in-vitro model. We used the Caco-2/TC7 cell line, a clonal population of human colon carcinoma-derived Caco-2 cells that closely reproduces most of the morphological and functional characteristics of normal human absorptive enterocytes^42,43^. Initially, we ensured that neither the Caco-2/TC7 basal culture medium (**Figure S1A**) nor the 1% Triton X-100, used for preparing the cell lysates (**Figure S1B**) directly affected phage viability. T4, M13, and ΦX174 phages, along with the fluorescent tracer FD4, were introduced into the apical compartment at the beginning of the experiment. A scheme of the experimental setup is shown in **Figure 2A**.

**Figure 2:**
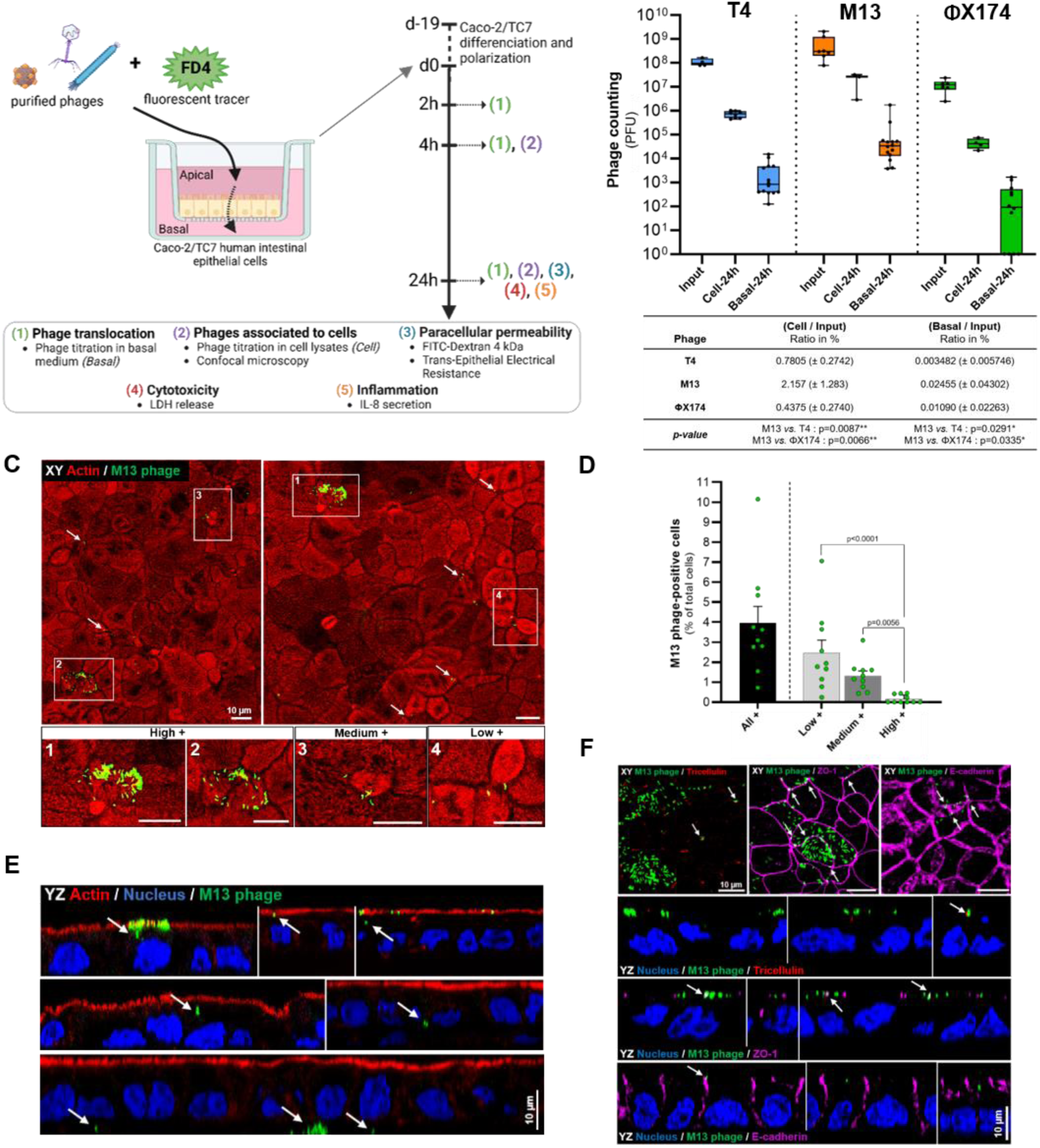
Internalization and translocation of phages across human intestinal epithelial cells. **(A)** Overview of the experimental setup. **(B)** Titration of T4, M13 and ΦX174 phages associated to Caco-2/TC7 cells *(cell)* and translocated through the monolayer *(basal)* after 24 h of incubation. The input refers to the phages initially added to the apical compartment. A minimum of four independent experiments [n ≥ 4] were performed. The table displays the percentage of phages associated to cells and translocated across Caco-2/TC7, normalized to the input. For details on quantification and cut-off criteria, see the Method section. **(C)** Images of CF488-labeled M13 phages interacting with Caco-2/TC7 cells following a 24 h of incubation. Examples of High+, Medium+ and Low+ fields are shown in insets, corresponding to different fluorescence levels of Caco-2/TC7 positive for CF488-labeled M13 phages. **(D)** Quantification of the percentage of Caco-2/TC7 ceils positive for CF488-labeled Ml3 phages, as shown in panel (C). Cells were categorized based on the fluorescence intensity of phages from analysis of 10 randomly selected images. **(E)** YZ views of CF488-labeled M13 with Caco-2/TC7 cells following 24 h of incubation. **(F)** Images (XY and YZ views) of CF488-labeled M13 phages combined with immunofluorescence stainings of tricellulin (tricellular tight junction), Zonula Occludens-1 (ZO-1, tight junction) and E-cadherin (adherens junction) after 24 h of incubation. In all fluorescence images, M13 phages are covalently conjugated with CF488 fluorophore (green), actin is labeled with Alexa A546-phalloidin (red), and nuclei with DAPI (blue). White arrows indicate phages colocalizing with actin (C), Tricellulin (F), ZO-1 (F), and E-cadherin (F) or internalized within cells (E). Scale bar, 10 pm. Data are represented as mean ± SEM.

After 2 h of incubation, only M13 phages were detected in the basal medium (data not shown). However, after 4 h, all three phages were recovered in the basal medium, with the rate of M13 translocation being significantly higher than that of T4 (**Figure S2**). After 24 h, phage translocation increased approximately 10-fold relative to that after 4 h. Depending on the phage, 0.0035% to 0.02% successfully translocated across the monolayer and between 0.44% and 2.2% were found associated with intestinal epithelial cells (**Figure 2B**). Interestingly, M13 phages showed significantly higher levels of association with the cells and translocation than the other two phages (**Figure 2B**) suggesting a phage-dependent translocation process.

We explored the localization of M13 phages to better understand how they translocate through intestinal epithelial cells by covalently conjugating them with CF488 fluorophores, and following them by confocal microscopy. After 24 h of incubation, M13 phage particles were colocalized with actin of the microvilli, both as aggregates and isolated phages, leading to an average of 4% of cells positive for CF488-labelling (**Figure 2C-D**). Large aggregates of M13 phage particles on intestinal epithelial cells were 16-fold less frequent than isolated phages (**Figure 2D**). Examination along the YZ axis confirmed that M13 phage particles were located at the level of microvilli which were revealed by actin staining with a weak proportion internalized within intestinal epithelial cells (**Figure 2E**). They were also visualized below the epithelial monolayer, in accordance with our titration of phages in the basal medium, demonstrating that functional phages are able to cross intestinal epithelial cells (**Figure 2E**). We then examined whether M13 phages could also translocate through the intestinal epithelium using paracellular pathways. We labelled tight and adherens junction proteins and observed colocalizations between M13 phage particles and the junction proteins, tricellulin, ZO-1, and E-cadherin, suggesting that phages can also be associated with intercellular junctions although such subcellular localization was not predominant (**Figure 2F**).

We assessed whether phages are damaged during translocation across the epithelial barrier by quantifying T4 phage particles both by plaque assays and qPCR. There were no differences between PFU counts and quantification by qPCR, indicating that most translocated phages remain functional (**Figure S3**).

No differences in epithelial barrier integrity were observed between Caco-2/TC7 cells incubated with or without phages (CTL), as measured by FD4 flux, trans-epithelial electrical resistance (TEER), lactate dehydrogenase release, and IL-8 pro-inflammatory chemokine secretion (**Figure S4A-D**). These results indicate that phages do not themselves induce hyperpermeability, cytotoxicity, or inflammation.

These findings provide evidence that under physiological conditions, all three studied phages are able to translocate across intestinal epithelial cells without disrupting the barrier function, with M13 phages exhibiting a higher translocation capacity.

### Phages internalize by endocytosis and translocate across endothelial cells without disrupting barrier integrity

Next, we investigated the fate of phages after their translocation across the epithelial barrier, hypothesizing that they might reach the bloodstream by crossing the intestinal blood vessels. As the endothelial barrier constitutes the lining of intestinal capillaries, we focused on how phages interact with this barrier.

HUVECs were cultured on a permeable membrane to establish a confluent monolayer. T4, M13, or ΦX174 phages, along with the fluorescent tracer FD4, were introduced into the apical compartment (**Figure 3A**). We first ensured that the HUVECs basal culture medium did not directly affect phage viability (**Figure S1C**). All three phages, T4, M13, and ΦX174, showed a high level of translocation across HUVECs within 1 h. Notably, only ΦX174 phages showed a significant further increase after 24 h, and no significant difference was observed between the three phages (**Figure 3B**). The translocation rate of T4, M13, and ΦX174 across the endothelial barrier was 346-fold, 285-fold and 806-fold higher, respectively, than that across the epithelial barrier (**Figure 2B and 3B**). These results indicate that phages can cross an endothelial barrier, regardless of the phage type and thus may be able to reach the bloodstream.

**Figure 3:**
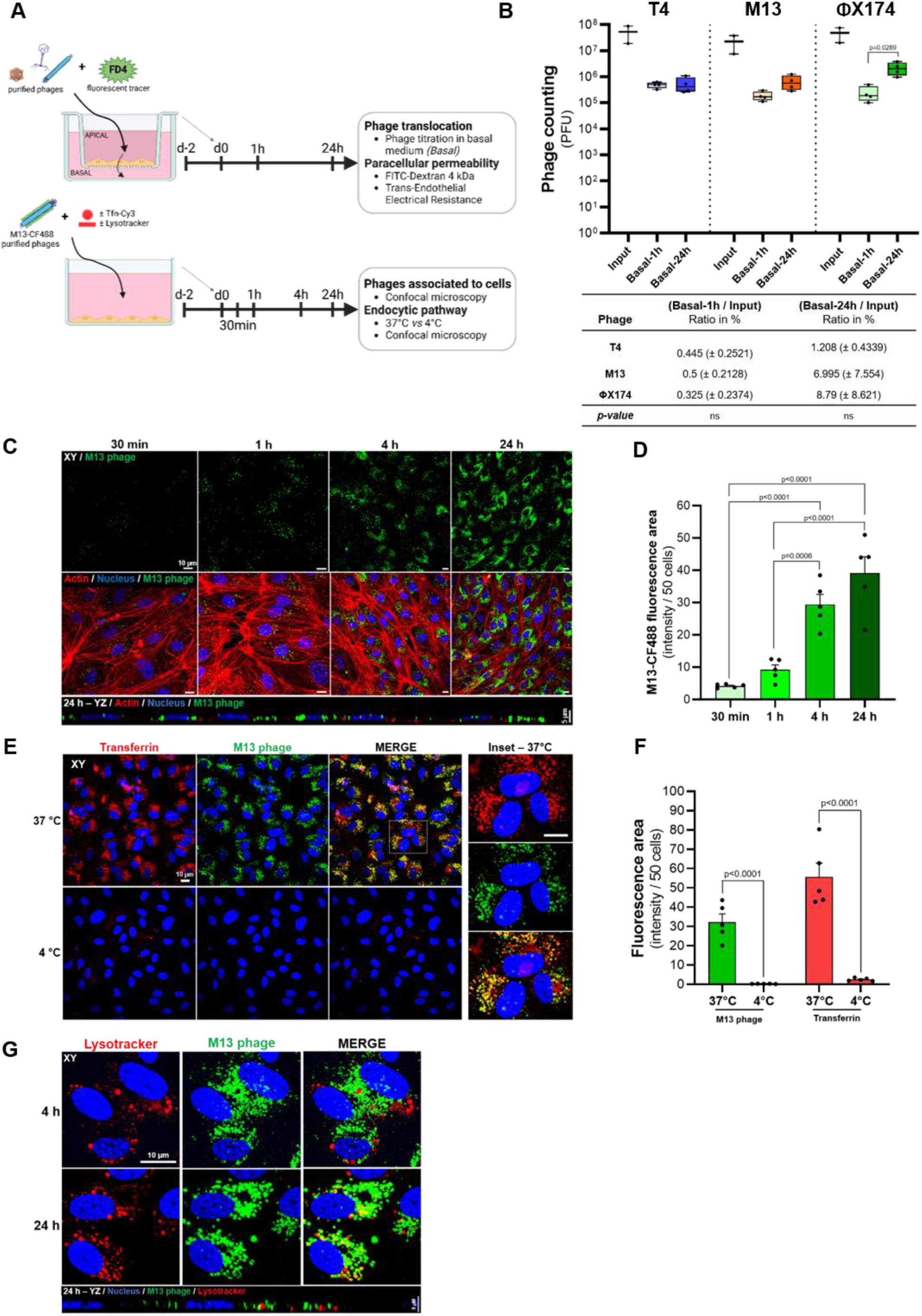
Internalization of phages through endocytosis and translocation across human endothelial cells. **(A)** Overview of the experimental setup. **(B)** Titration of T4, M13, and ΦX174 in the basal compartment *(basal)* after 1 h and 24 h of incubation. The input refers to the phages initially added to the apical compartment. The table displays the percentage of translocated phages across HUVECs, normalized to the input. **(C)** Images of CF488-labeled M13 phages with HUVECs cells after 30 min, 1 h, 4 h, and 24 h of incubation. Actin was labeled with Alexa A546-phalloidin (red), and nuclei with DAPI (blue). The YZ view corresponds to 24 h of incubation. **(D)** Quantification of the M13-CF488 fluorescence shown in panel (C). Data represent mean values ± SEM obtained from the analysis of 5 randomly selected images. **(E)** Images of CF4884abeled M13 phages, and Cy3-labeled transferrin in HUVECs cells after 4 h, at 37°C, and 4°C. **(F)** Quantification of the red fluorescence area (transferrin), and green fluorescence area (M13 phages) in HUVECs following 4 h, at 37°C, and 4°C, shown in panel (E). Data represent mean values ± SEM obtained from analysis of 5 randomly selected images. (G) Images of CF488-labeled M13 phages, and Lysotracker Deep Red in HUVECs cells after 4 h, and 24 h. The YZ view corresponds to 24 h of incubation. Scale bar, 10 pm.

We then investigated the mechanism by which phages translocate across the endothelial barrier, hypothesizing that they are internalized by endothelial cells through endocytosis. We introduced CF488-labelled M13 phage particles (**Figure 3A**) and observed significant accumulation within HUVECs from 30 min to 24 h, with perinuclear localization (**Figure 3C-D**), showing that HUVECs are highly permeable to phages. We compared the intracellular trafficking of fluorescent M13 phage particles with that of Cy3-labelled transferrin, used as an endocytosis cargo control^75^, at both 37°C and 4°C (a temperature known to inhibit endocytosis). M13 phage particles and Cy3-labelled transferrin were simultaneously introduced onto a HUVECs monolayer (**Figure 3A**). We observed a strong fluorescent transferrin signal within the HUVECs from 30 min up to 4 h of incubation at 37°C, confirming, as expected, its internalization by endocytosis (**Figure S5A-B and 3E-F**). By contrast, after 30 min of incubation at 37°C, the fluorescent M13 phage signal was weak (**Figure S5A-C**), indicating that transferrin appears to be internalized more rapidly than M13 phages within HUVECs. After 1 to 4 h of incubation, robust colocalization was observed between transferrin and M13 phage particles, suggesting their convergence within the endocytic pathway (**Figure S5A and 3E**). Importantly, both intracellular transferrin and M13 phage signals were much weaker at 4°C than 37°C (**Figure S5A-C and Figure 3E-F**), indicating that M13 phages are internalized through an active endocytic process. No significant colocalization was detected over time between M13 phage particles and lysotracker, which labels acidic compartments (**Figure 3G and S5D**), suggesting that most phages evade lysosomal degradation.

Furthermore, none of the tested phages modified the permeability of the endothelial barrier by themselves, assessed by xCELLigence, FD4 passage or TEER **(Figure S6A-D)**. Overall, these results show that phages can easily translocate across an endothelial barrier through endocytosis and without inducing deleterious effects on the epithelium, thus demonstrating their potential to efficiently reach the bloodstream.

### A compromised intestinal barrier facilitates ΦX174 translocation but has no impact on M13 or T4 translocation

CD is associated with a defect in the intestinal barrier, notably characterized by increased intestinal permeability^2,17,25^.. We explored whether such hyperpermeability could lead to higher phage translocation by mimicking alterations of intestinal epithelial and endothelial barriers in-vitro.

First, Caco-2/TC7 cells were treated with either EGTA to globally disorganize cell-cell junctions or with pro-inflammatory cytokines to trigger an inflammatory condition, before introducing phages (**Figure 4A**). As expected, IFNγ + TNFα and EGTA altered epithelial barrier function (**Figure S7A-B**) by significantly increasing paracellular permeability to both ions and macromolecules relative to untreated cells, with EGTA having a more pronounced effect than the cytokines (**Figure S7A-B**). IFNγ + TNFα treatment also led to a significant increase in cytotoxicity relative to the control and EGTA conditions, and both treatments increased IL-8 secretion (**Figure S7C-D**). EGTA treatment significantly increased the translocation of ΦX174 phages relative to the control, and IFNγ + TNFα conditions, whereas the translocation of M13 and T4 was not modified by these treatments (**Figure 4B**). Furthermore, there was a significant correlation between the level of paracellular permeability to both ions and macromolecules and the translocation of ΦX174 phages, but no significant correlation was observed for T4 and M13 phages (**Figure 4D and Table 2**). Similar to the physiological context, under compromised barrier conditions, there were no differences between PFU counts and quantification by qPCR, suggesting that most translocated phages remain functional and that barrier defects do not affect phage infectivity (**Figure S3**).

**Figure 4:**
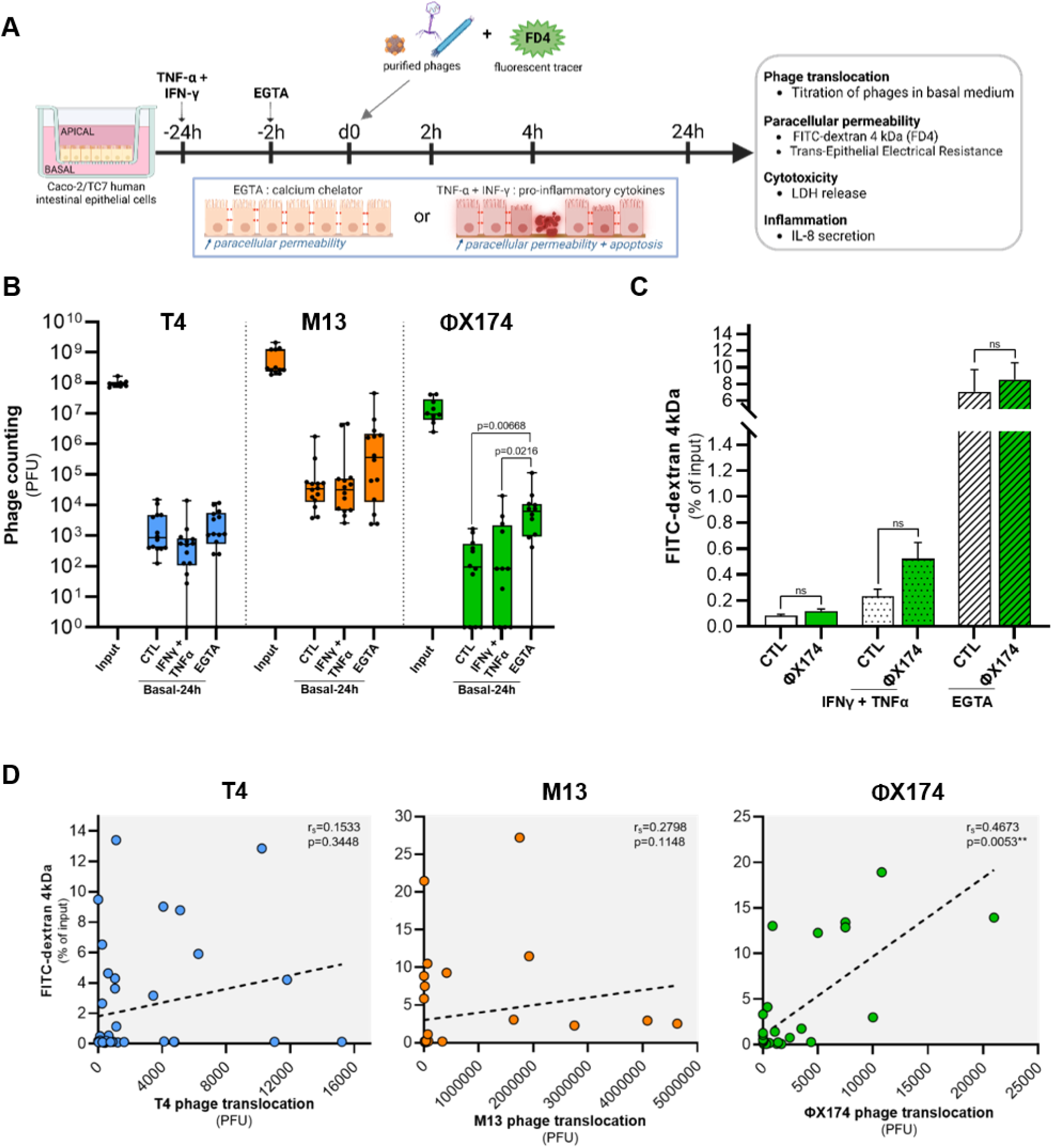
A compromised intestinal barrier facilitates ΦX174 translocation across intestinal epithelial cells. **(A)** Overview of the experimental setup. **(B)** Titration of T4, M13, and ΦX174 phages in the basal compartment (basal) after 24 h of incubation in the apical compartment. The input refers to the phages initially added to the apical compartment. Conditions include healthy barrier condition (CTL) relative to compromised barriers conditions induced by IFNγ and TNFct or EGTA. **(C)** FITC-Dextran 4 kDa (FD4) passage from the apical to the basal compartment during 24 h in presence of ΦX174 phages compared to control without phages (CTL) condition under healthy, and compromised barrier conditions (IFNγ + TNFα or EGTA). **(D)** Correlation analysis between the translocation rates of T4, M13, or ΦX174 phages in all conditions (CTL, EGTA, and IFNγ + TNFα) and FD4 flux. Spearman’s test was used to assess the correlation between the two variables tested. For all experiments, a minimum of four independent experiments [n ≥ 4] were performed. Data are represented as mean ± SEM.

**Table 2.**
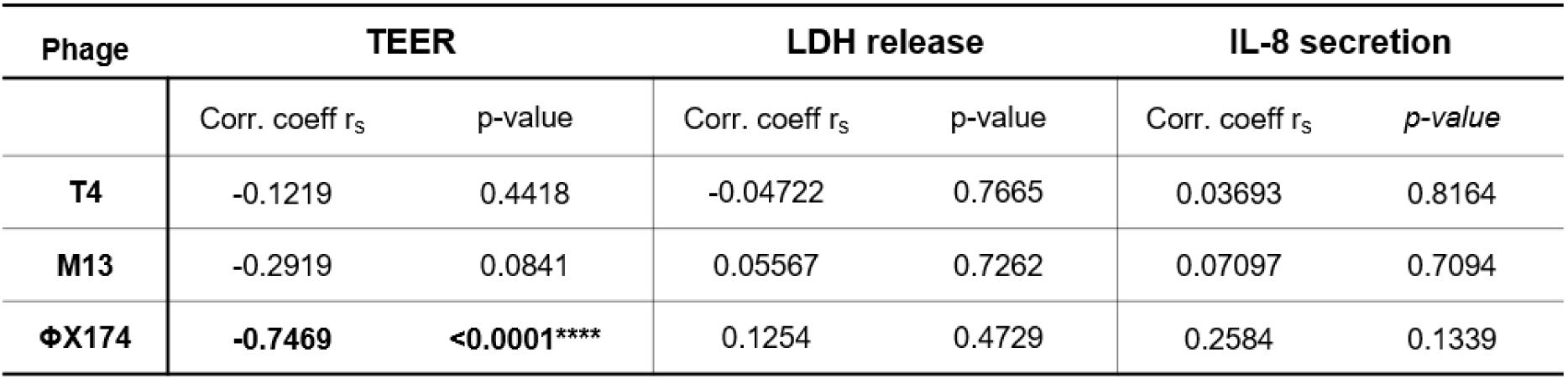
Correlations between phage translocation and TEER, LDH release, or IL-8 secretion in Caco-2/TC7 intestinal epithelial cells. Correlation analysis were performed between the translocation rates of T4, M13, or (ΦX174 phages in all conditions (CTL, EGTA, and IFNγ + TNFα) and TEER, LDH release, or IL-8 secretion. Spearman’s test was used to assess the correlation between the two variables tested.

In addition, in Caco-2/TC7 cells under compromised barrier conditions, ΦX174 phages did not induce hyperpermeability (**Figure 4C and Table S4**), cytotoxicity (**Figure S8E**), or exacerbate inflammation (**Figure S8H**). Similar results were observed for T4 (**Figure S8A-C-F),** and M13 phages (**Figure S8B-D-G)**, indicating that when barrier integrity is compromised, phages do not trigger further deleterious effects upon translocation.

We then assessed the effects of an endothelial barrier defect on phage translocation. HUVECs cultured on permeable membranes were incubated with IFNγ and TNFα to mimic an inflammatory condition. Such treatment resulted in a significant increase in FD4 flux and a decrease in TEER (**Figure S9A-B**), without inducing LDH release (**Figure S9C**). These results demonstrate that, as expected, pro-inflammatory cytokines increase paracellular permeability of the endothelial barrier without inducing cytotoxicity. The inflammatory state did not affect the translocation rate of T4, M13, or ΦX174 phages across HUVECs (**Figure S9D-F**), suggesting that phages do not mainly translocate across the endothelial barrier through the paracellular pathway.

Overall, these findings show that inflammatory conditions have no impact on phage translocation through the endothelial barrier but facilitate the translocation of ΦX174 phages across the epithelium.

### CD patients exhibit a signal of higher translocation of phages from the gut to the blood, including those of the *Microviridae* family

After demonstrating that phages can translocate under physiological and compromised barrier conditions, we sought to validate these findings in a cohort of patients with CD, and examined whether this pathological condition of gut barrier deficiency leads to higher phage translocation from the intestinal lumen to the bloodstream. Blood and stool samples were collected from 15 CD patients and 14 HS (**Table S1**) and their viral metagenome was analyzed (**Figure 5A**). Among the viral operational taxonomic units (vOTUs) identified across the 29 participants, 14,843 were found exclusively in stool, whereas 1,746 were unique to blood and 48 (0.3% of total vOTUs) were detected in both stool and blood samples, regardless of whether they came from the same participant (**Figure S10**). A complete analysis of these samples is reported in a separate study from our laboratory and revealed significant differences in the beta-diversity of the blood virome of CD patients relative to HS^32^.

**Figure 5:**
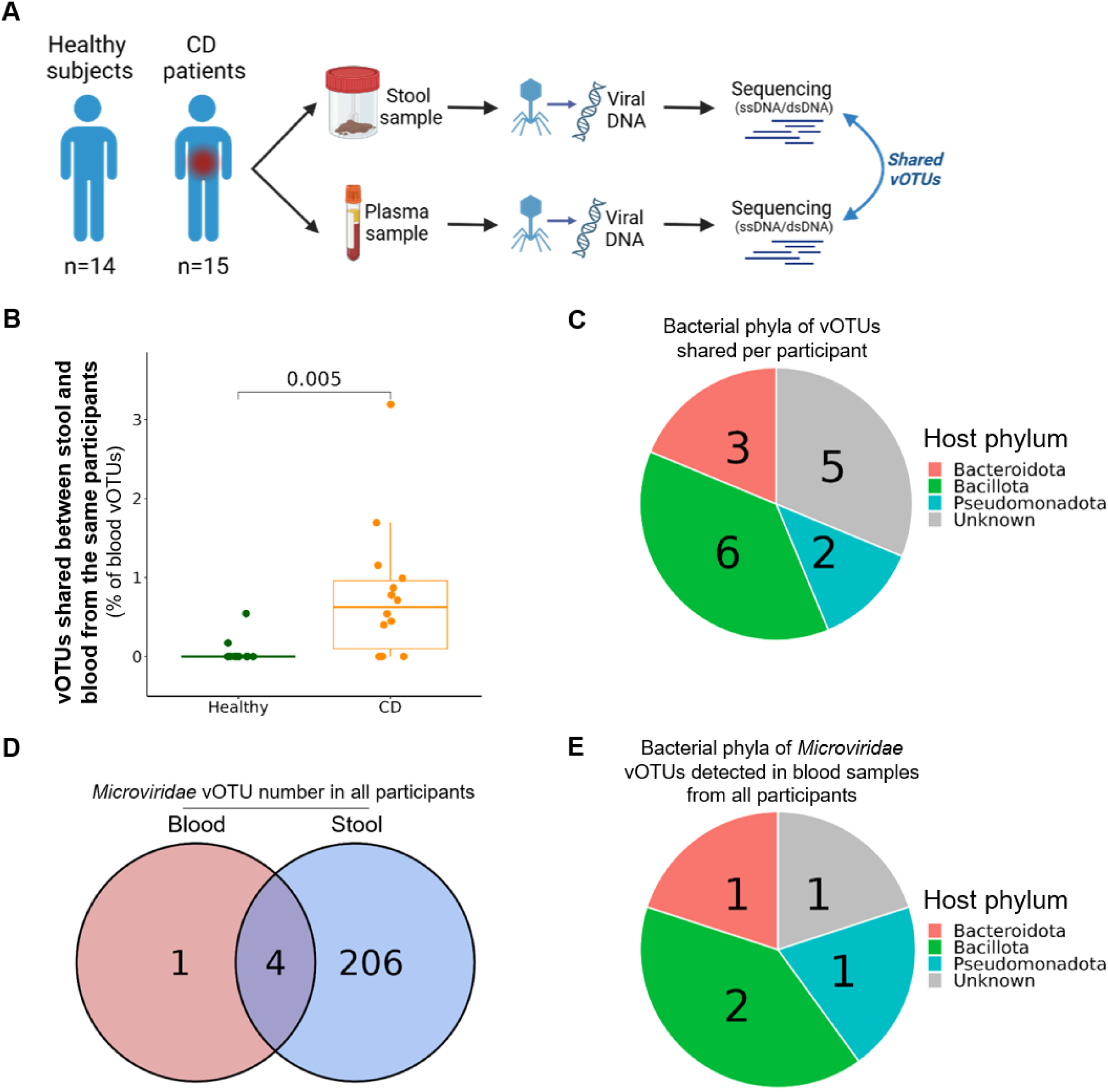
Analysis of shared stool and blood viromes in healthy subjects and CD patients. **(A)** Overview of the cohort sample collection, and experimental setup. **(B)** Percentage of unique vOTUs shared between stool and blood from the same participants in CD patients and healthy subjects. **(C)** Distribution of predicted bacterial host phyla for the 16 vOTUs shared between stool and blood samples from the same participants. **(D)** Venn diagram illustrating the overlap of *Microviridae* vOTUs identified in blood and stool samples across **the** 29 participants. **(E)** Distribution of predicted bacterial host phyla for the *Microviridae* vOTUs detected in blood samples from the 29 participants.

Here, to investigate the role of the gut barrier, we focused on vOTUs shared between stool and blood samples for each of the 29 participants, reflecting the level of phage transfer (**Figure 5A**). We found a significantly higher number of vOTUs shared in CD patients than HS, indicative of a potential increase in phages translocating from the intestinal lumen to the bloodstream (**Figure 5B**). Since most of the vOTUs correspond to phages, we looked for their bacterial hosts. Among the 16 vOTUs shared between blood and stool samples per participant (from all the CD and HS), six infected hosts belonging to the phylum Bacillota (previously known as Firmicutes) and three infected Bacteroidota (previously Bacteroidetes), which are the two most abundant phyla in the human gut microbiota^76^, suggesting a gut origin for these vOTUs (**Figure 5C**). These findings support the hypothesis that phage translocation from the intestinal lumen to the bloodstream is increased in CD patients.

Given our previous observation that ΦX174 phage translocation is higher when the intestinal barrier is impaired in the Caco-2/TC7 epithelial cell model, we postulated that *Microviridae* phages may be more likely to cross the gut and reach the bloodstream. From the total vOTUs dataset (blood and stool), 211 *Microviridae* vOTUs were identified, accounting for 1.3% of vOTUs (**Figure 5D**). All 29 participants harbored *Microviridae* in their stool samples, totaling 210 vOTUs (**Figure 5D**). Notably, there was no difference in *Microviridae* diversity or relative abundance between CD and HS in stool samples (**Figure S11**). In blood samples from the 29 participants, only 5 *Microviridae* vOTUs were found across two CD and one HS, with 4 vOTUs detected in both stool and blood samples (**Figure 5D**). This overlap shows that most *Microviridae* found in blood are also present in stool. Moreover, these 5 blood *Microviridae* vOTUs infected hosts belonging to Bacillota and Bacteroidota bacterial phyla (**Figure 5E**), suggesting a gut origin of the blood *Microviridae*.

In addition, among the 16 vOTUs shared between stool and blood samples from each participant (from CD and HS), above mentioned, two belong to *Microviridae*, representing approximately 13% of the transferred vOTUs from this family. These two *Microviridae* vOTUs originated from the same CD patient and showed a strong gut association (>90% average nucleotide identity homology against gut virus databases; see Methods). Despite the low number, this substantial enrichment (10X) relative to the 1.3% prevalence of *Microviridae* in our total dataset of 16,637 vOTUs suggests that phages from the *Microviridae* family may more frequently cross the intestinal barrier than other phage families and preferentially reach the bloodstream.

Overall, virome analysis confirms that phages are able to translocate from the gut to the blood, particularly *Microviridae* phages, and indicates that translocation is more frequent in CD patients, affected by impaired gut barrier function.

## DISCUSSION

Although a few studies have recently highlighted the interaction of phages with various cell types33–38, such interactions are still poorly understood in both physiological and disease contexts.

Despite the tightly regulated nature of the intestinal epithelial barrier, we observed that phages are able to translocate across intestinal epithelial cells, with different rates between the tested phages. In this vein, a recent study reported minimal translocation of phages targeting *E. coli* across intestinal epithelial cells^34^ and the translocation process appeared to be size-dependent^33,35^. Our study is the first to simultaneously compare phages with such distinct morphologies, *i.e. Inoviridae* filamentous phages, caudated *Straboviridae* phages, and *Microviridae* phages of smaller size. In the physiological context, we observed a higher level of translocation for M13, a filamentous phage, suggesting a better translocation capacity for *Inoviridae* than *Straboviridae* and *Microviridae*. Fluorescently tagged phages were observed at the microvilli, internalized within and crossing intestinal epithelial cells. Although rare, we noted aggregates of M13 phages on intestinal epithelial cells, a phenomenon not observed on endothelial cells. The mechanisms underlying such aggregation is unknown but possibilities include a specific cellular signal that trigger phage aggregation or trapping within the microvilli. In addition, an active process induced by phages is possible, as M13 phages have been shown to increase gene expression when interacting with specific human cells^77^. Phage translocation across human cells is suggested to be vectorized from the apical to the basal side^35,78^ and, based on microscopy images, we suggest that microvilli could serve as gateways for phages to internalize via transcytosis. For example, phages might interact with cell surface proteins that share structural similarities with bacterial phage receptors^38^. Another possibility is the potential for phages such as M13, which are approximately six nanometers in diameter^79^, to use paracellular pathways. We found the phages to be rarely associated with tight and adherens junctions. Although our current data do not allow a definitive answer, we hypothesize that M13 phages may primarily use the transcytosis pathway, as demonstrated in other cell lines^80^.

Our study demonstrates that phages interact with the endothelial barrier in a markedly different way than with the epithelial barrier. Endothelial cells were over 500 times more permeable to phages, and this translocation occured independently of phage type. Highly polarized epithelial cells, which form a selective barrier with tightly organized junctions, contrast sharply with gut vascular endothelial cells, which have fewer and more loosely organized tight junctions, leading to greater permeability^3^. Supporting our findings, small macromolecules, such as those of approximately 4 kDa, can easily diffuse through the endothelial barrier into the bloodstream^81^, whereas intestinal epithelial cells restrict their passage^82,83^. We also found that M13 phages are internalized by endothelial cells through endocytosis, a process confirmed by other studies in different cell types^33,35,38,80^. Although phage internalization in the endothelial barrier appears to be active, the exact endocytic pathway involved is still unclear^84^. According to Tian et al., the size of M13 filamentous phages may be compatible with macropinocytosis^80^, but T4 and ΦX174 phages could use different endocytic pathways^76^.

Pharmacokinetic studies have detected phages in the bloodstream of mice, highlighting their ability to cross biological barriers^85–87^. Our study provides a detailed explanation of this process, demonstrating that functional phages crossing the epithelial barrier can easily reach the bloodstream due to the high permeability of the endothelial barrier. These findings are consolidated by our ex-vivo experiments and a parallel study from our team indicating that approximately 20% of phages found in the bloodstream likely originate from the gut^32^. In addition, phages may translocate through lymphatic vessels, allowing them to access the lymphatic system, reach the bloodstream and potentially spread to distant organs.

Using in-vitro models, we observed that the impairment of epithelial barrier function, a hallmark of CD^2,16,17,25^, leads to increased translocation of ΦX174 phages, a member of the *Microviridae* family. This increase in phage translocation directly correlates with the level of paracellular permeability, emphasizing that *Microviridae* phage translocation is particularly sensitive to the integrity of the intestinal epithelial barrier. We speculate that phage size may also affect translocation rate, as ΦX174 is the smallest of the three phages tested in this study. In addition, we observed that CD patients exhibit a higher number of viral sequences shared between stool and blood samples per participant than healthy subjects, indicating increased phage translocation from the gut to the bloodstream when the intestinal barrier is compromised. Although early studies suggested changes in the gut virome of CD patients, recent analyses have not detected a CD-specific pattern due to the high variability of the gut virome^31^. Despite such variability, viromes of IBD patients were found to exacerbate inflammation *in vivo*^36,88,89^. Our virome analysis showed the blood phageome to be enriched in *Microviridae* phages, which are also a large community of the gut virome^26^, suggesting that they are preferentially translocated from the gut to the blood. The pattern of phage translocation in CD patients may contribute to a unique blood viral community, but whether it is involved in CD pathogenesis, is yet to be determined.

Another key consideration is the direct impact of phages on human cells, especially concerning their use as therapeutic agents, known as phage therapy. Certain phages have been identified to be highly immunogenic^36,90,91^ and can increase endothelial cell permeability^92^, whereas others elicit a minimal immunological response^93,94^ and do not trigger pro-inflammatory reactions^95^. Our data show that T4, M13, and ΦX174 functional phages can translocate without compromising the function of the epithelial or endothelial barriers in both physiological and inflammatory contexts, supporting their safety in phage therapy. This evidence also strengthens the therapeutic potential of retargeted M13 phages, recently shown to penetrate cancer cells^96^ and tumor 3D spheroids^97^. Furthermore, in the context of IBD, we found that only a small fraction of phages can cross the intestinal epithelium, which is a highly selective barrier, suggesting that the effectiveness of phage therapy in the intestinal lumen may not be significantly affected by the translocation rate. The effect of phages on human cells could have important implications for health, but further studies are needed to validate these findings, particularly with intestinal phages.

In conclusion, our results highlight how distinct phages can cross the intestinal barrier under both physiological and inflammatory conditions, with translocation influenced by phage morphology, size, and barrier integrity. This study supports the potential safety of phages as therapeutic agents, strengthens our understanding of phage dynamics in CD and pave the way for exploring phage translocation in other disorders characterized by increased intestinal permeability.

## ACKNOWLEDGEMENTS

We thank Marie-Agnès Petit et Romain Sausset from the Phage Team at the Micalis Institute, Inrae Jouy-en-Josas, for providing the phages and bacterial strains, Juliane Selot from the "TGFb and cancer" team of the CRSA for supplying Cy3 labeled-transferrin, and Dr. Jessica Dahan Saal from Bluets Maternity Hospital (Pierre Rouquès Hospital, Paris) for the generous donation of umbilical cords. We also thank Marie-Agnès Petit, Etienne Morel, and Cédric Delevoye for valuable discussion as well as Raphaëlle Liquard and Lola Savouré for their assistance with human cell care. We are grateful to Tatiana Ledent and the staff in charge of animal housing and care at the animal care facility at the CRSA. In addition, we also thank the CISA platform at the CRSA, particularly Romain Morichon and Annie Munier for their support, made possible with financial support from Sorbonne Université and from ITMO Cancer of Aviesan and INCa on funds administered by Inserm. For virome sequencing, we thank M. Monot, L. Ma, and L. Lemée from the Biomics Platform, C2RT, Institut Pasteur, Paris, France, supported by France Génomique (ANR-10-INBS-09) and IBISA. Finally, we acknowledge Biorender which was used to create the figures of the experimental setup. This research was supported by grants from the French National Research Agency (ANR), including VENTRIS (n°ANR-21-CE14-0019-01) and PRIMAVERA (n°ANR-19-CE18-0028-01), as well as institutional funding from the Institut National de la Santé et de la Recherche Médicale (INSERM), Sorbonne Université, and the École Pratique des Hautes Études.

## SUPPLEMENTARY FIGURES

**Figure S1:**
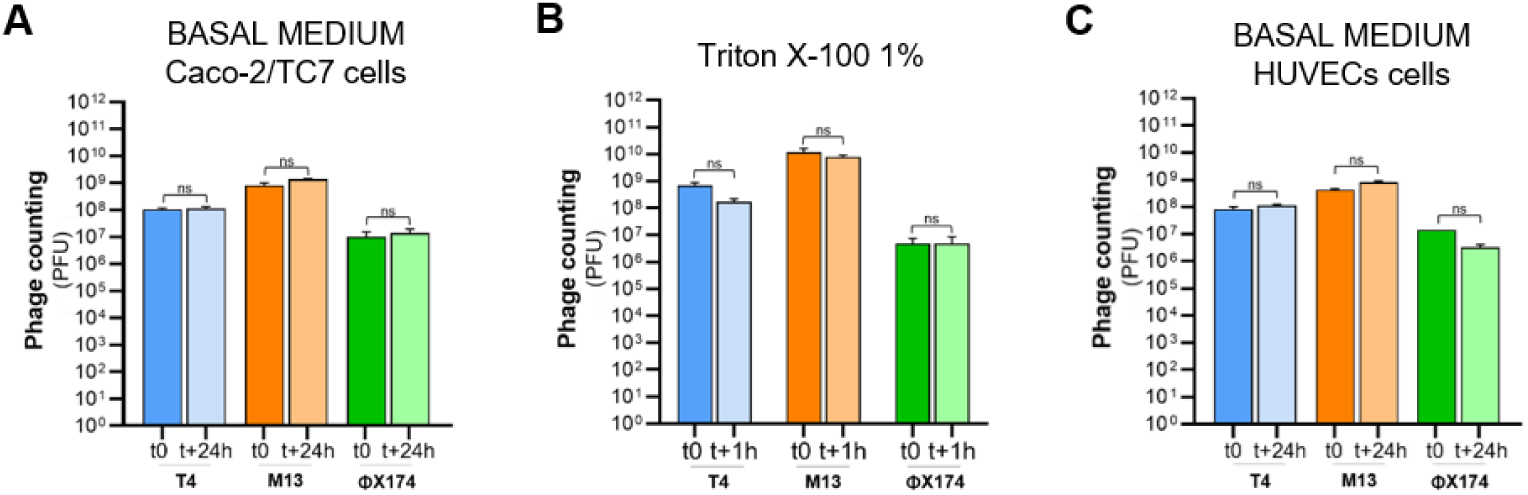
Cell culture medium and Triton X-100 do not impact phage viability. **(A)** Titration of T4, M13, and ΦX174 phages immediately after suspension (t0) and 24 h after incubation in Caco-2/TC7 culture medium. **(B)** Titration of T4, M13, and ΦX174 phages immediately after suspension (t0) and following a 1 h incubation with 1% Triton X-100 (t+1h), used in this study to lyse eukaryotic cells. **(C)** Titration of T4, M13, and ΦX174 phages immediately after suspension (t0) and 24 h after incubation in HUVEC culture medium (t+24h). All experiments were performed in triplicates [n = 3], Data are represented as mean ± SEM.

**Figure S2:**
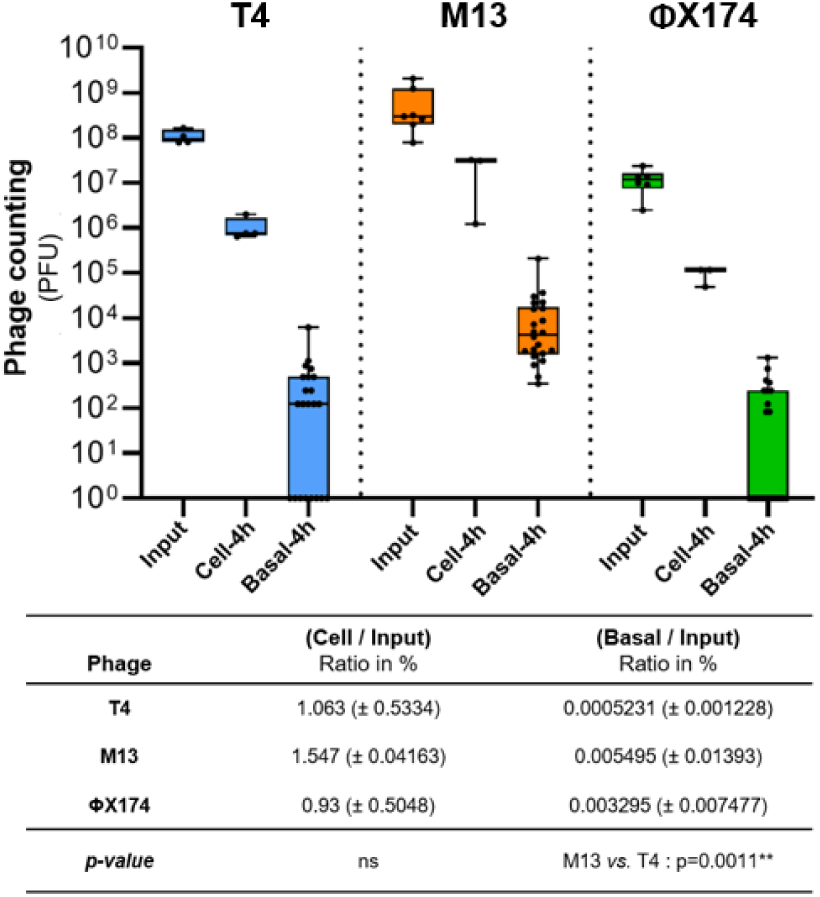
Internalization and translocation of phages across Caco-2/TC7 Intestinal epithelial cells after 4 h of inbubation. Titration of T4, M13, and ΦX174 phages associated to Caco-2/TC7 cells (cell) and translocated phages (basal) after 4 h of incubation in the apical compartment. The input refers to the phages initially added to the apical compartment. The table displays the percentage of phages associated to and translocated across Caco-2/TC7, normalized to the input. A minimum of four independent experiments [n ≥ 4] were performed. Data are represented as mean ± SEM.

**Figure S3:**
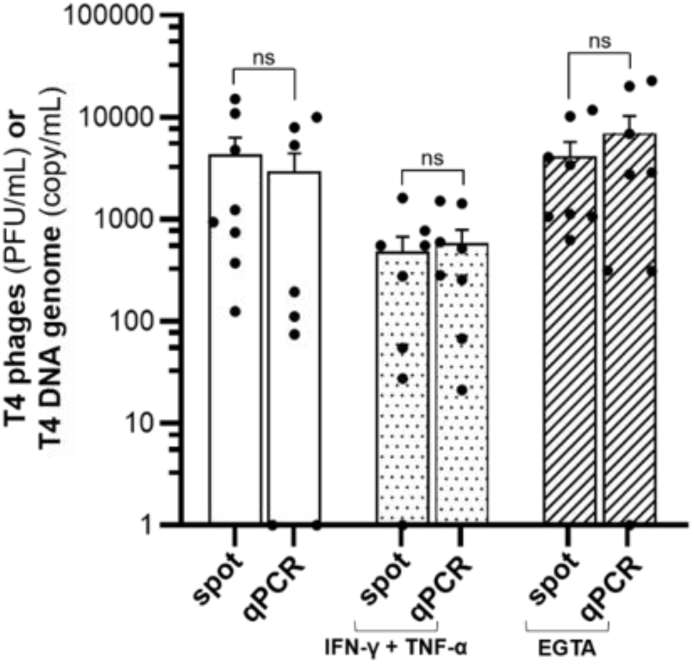
T4 phages that translocated across Caco-2/TC7 cells retain their functionality. Titration of T4 phages in the basal compartment after 24 h of apical incubation, measured by plaque assays (spot) and qPCR (a calibration curve using T4 DNA was used to determine the genome copy/mL) under both physiological, or compromised barrier conditions (IFN-γ + TNF-α, and EGTA). Two independent experiments were conducted in quadruplicate. Data are presented as mean ± SEM.

**Figure S4:**
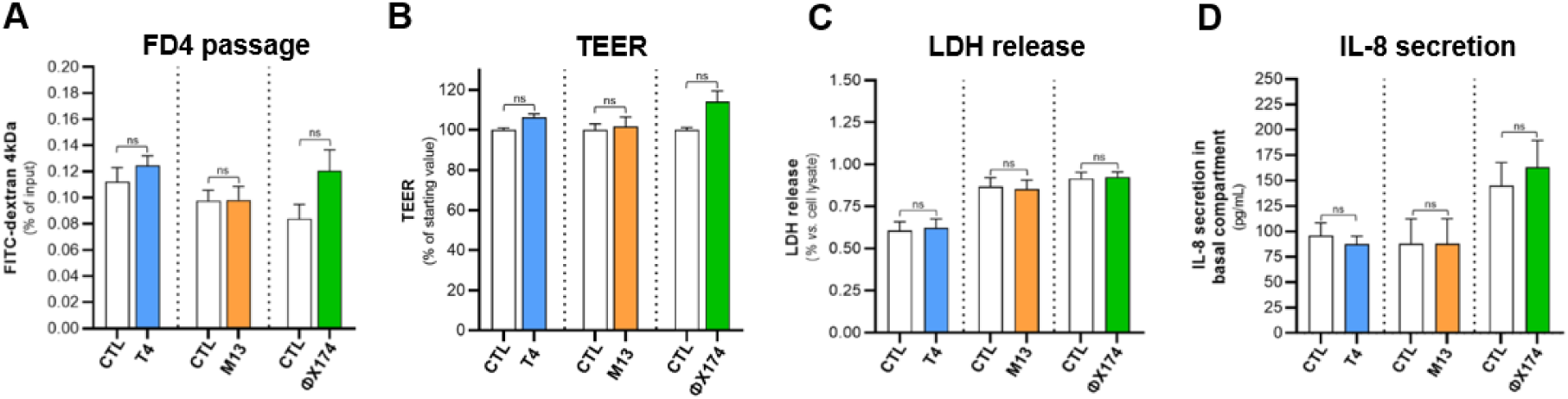
Phages do not induce hyperpermeability, cytotoxicity, or inflammation in Caco-2fTC7 cells under physiological condition. **(A)** FITC-Dextran 4 kDa (FD4) passage from the apical to the basal compartment during 24 h in the presence of the indicated phage compared to control (CTL) condition without phage. **(B)** Measure of transepithelial electrical resistance (TEER) after 24 h of incubation with medium containing no phages (CTL) or the indicated phages. Values are expressed as a percentage relative to the starting value for each well and then normalized to the control condition. **(C)** Release of lactate dehydrogenase (LDH) by the cells in the apical compartment, after 24 h of incubation with medium containing no phages (CTL) or the indicated phages. Values are expressed as the percentage of LDH measured in the whole cell lysate. **(D)** Secretion of pro-inflammatory cytokine IL-8 by the cells within the basal compartment after 24 h of incubation with medium containing no phages (CTL) or the indicated phages. For all the experiments, a minimum of four independent experiments [n ≥ 4] were performed. Data are represented as mean ± SEM.

**Figure S5:**
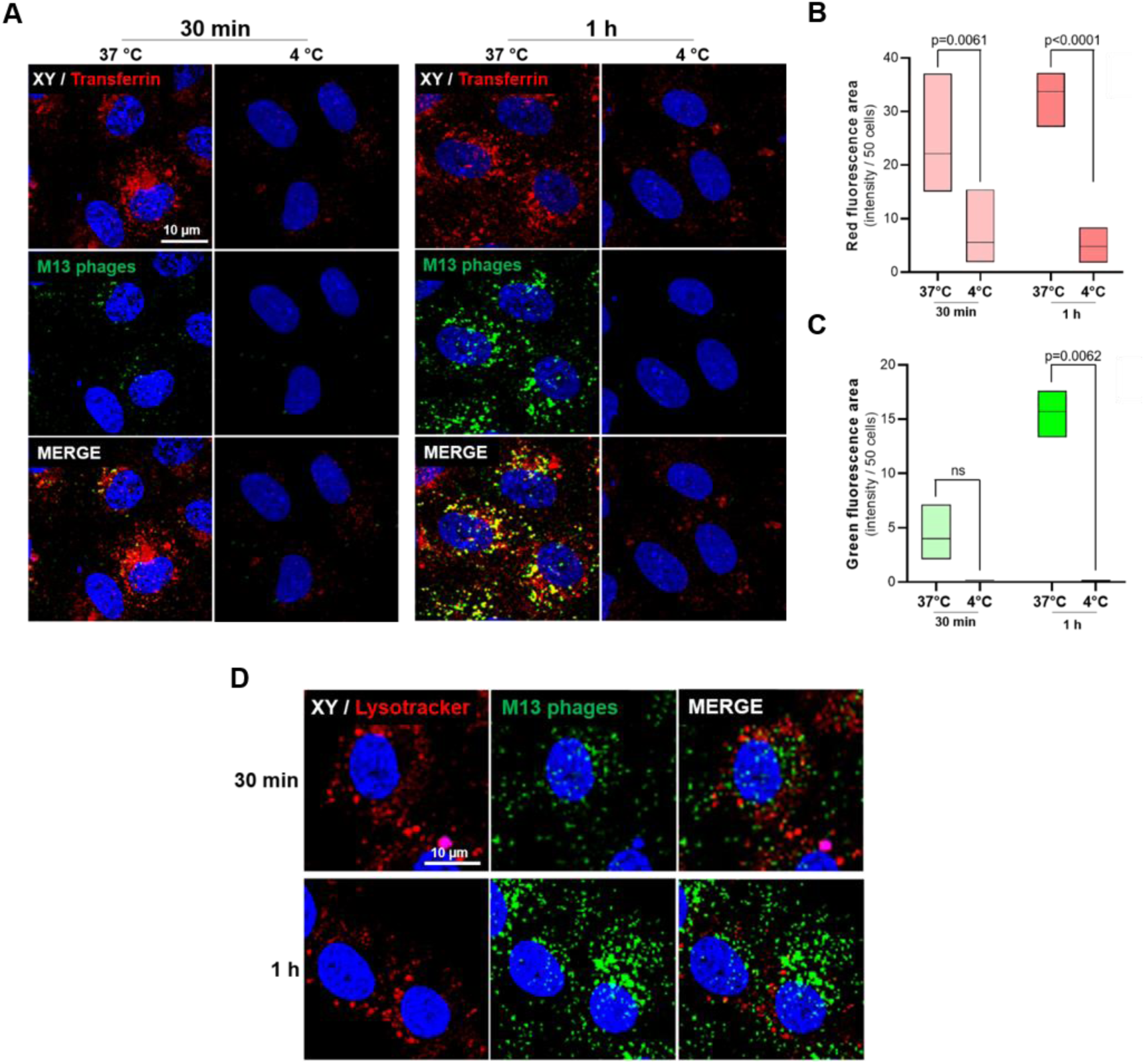
Visualization of phages in the endocytic pathway in HUVECs cells following short Incubation periods. **(A)** Images of CF488-labeled M13 phages, and Cy3-labeled transferrin in HUVECs after 30 min and 1 h, at 37°C, and 4°C. Quantification of the **(B)** red fluorescence area (transferrin), and **(C)** green fluorescence area (M13 phages) in HUVECs following 30 min and 1 h, at 37°C, and 4°C, as shown in panel (A). Data represent mean values obtained from analysis of 5 randomly selected images. b Images of CF488-labeled M13 phages, and Lysotracker Deep Red in HUVECs after 30 min and 1 h. Scale bar, 10 pm.

**Figure S6:**
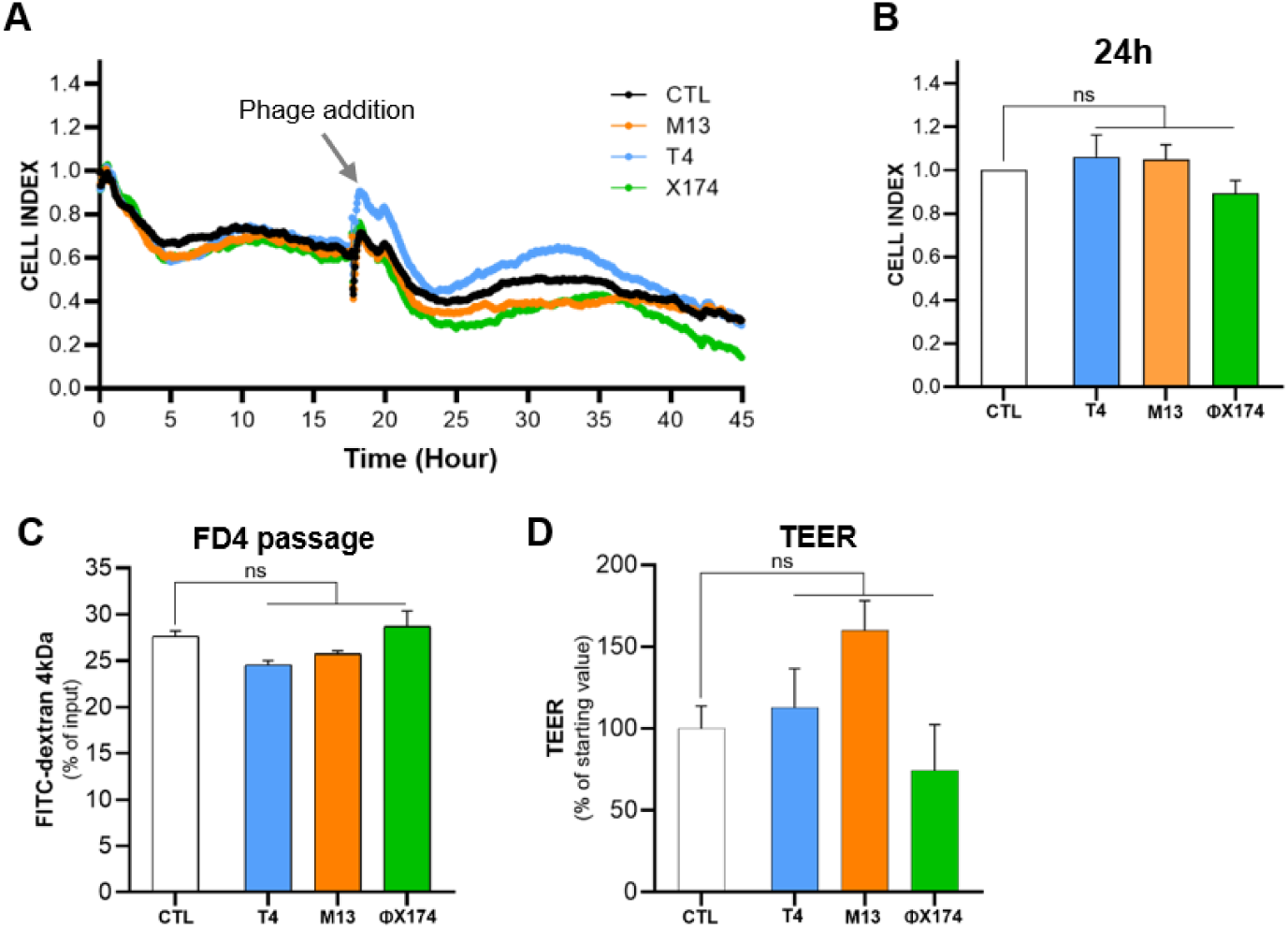
Phages do modify the permeability of HUVECs cells. **(A)** Representative plot of real-time permeability (cell index) of HUVECs in presence of the indicated phage relative to control (CTL) condition using xCELLigence system. **(B)** Cell index values obtained after 24 h in presence of the indicated phage relative to control (CTL) condition without phage after real time measure of HUVECs permeability, shown in panel (A). **(C)** FITC-Dextran 4 kDa (FD4) passage from the apical to the basal compartment after 24 h in presence of the indicated phage relative to control (CTL) condition. **(D)** Measure of transendothelial electrical resistance (TEER) after 24 h of incubation with medium containing no phages (CTL) or the indicated phages. Values are expressed as the percentage relative to the starting value for each well and then normalized to the control condition. For all these experiments, three independent experiments [n=3] were performed. Data are represented as mean ± SEM.

**Figure S7:**
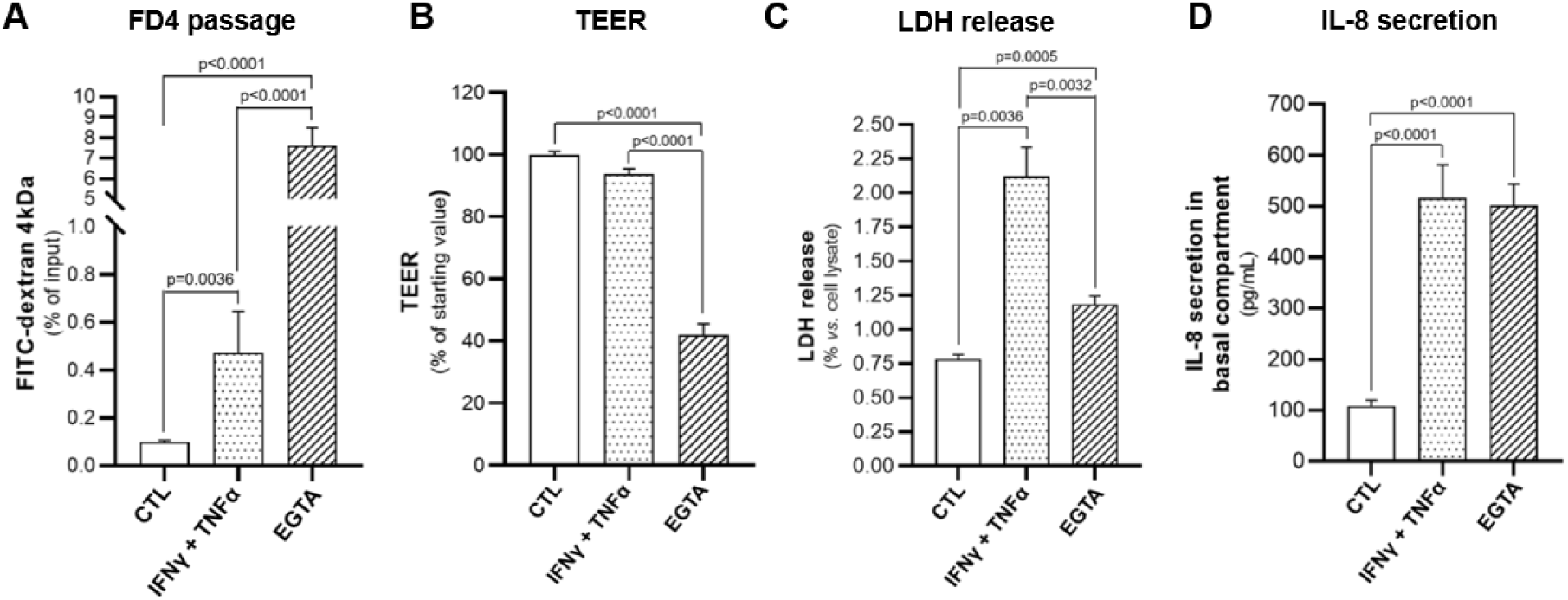
Pro-inflammatory cytokines, and EGTA induce a compromised barrier in Caco-2/TC7 cells. Caco-2/TC7 were treated with IFNγ + TNFα in the basal compartment during 48 h, or with EGTA in the apical compartment during 26 h. **(A)** FITC-Dextran 4 kDa (FD4) passage from the apical to the basal compartment (24 h of FD4 flux). **(B)** Measure of transepithelial electrical resistance (TEER) Values are expressed as a percentage relative to the starting value for each well and then normalized to the control condition. **(C)** Release of lactate dehydrogenase (LDH) from cells (expressed as the percentage of LDH measured in the whole cell lysate). **(D)** Secretion of pro-inflammatory cytokine IL-8 by the cells into the basal compartment. These data were pooled from control groups used in independent experiments involving T4, M13, and ΦX174 phages, with at least 3 independent experiments conducted for each [n ≥ 9], Data are represented as mean ± SEM.

**Figure S8:**
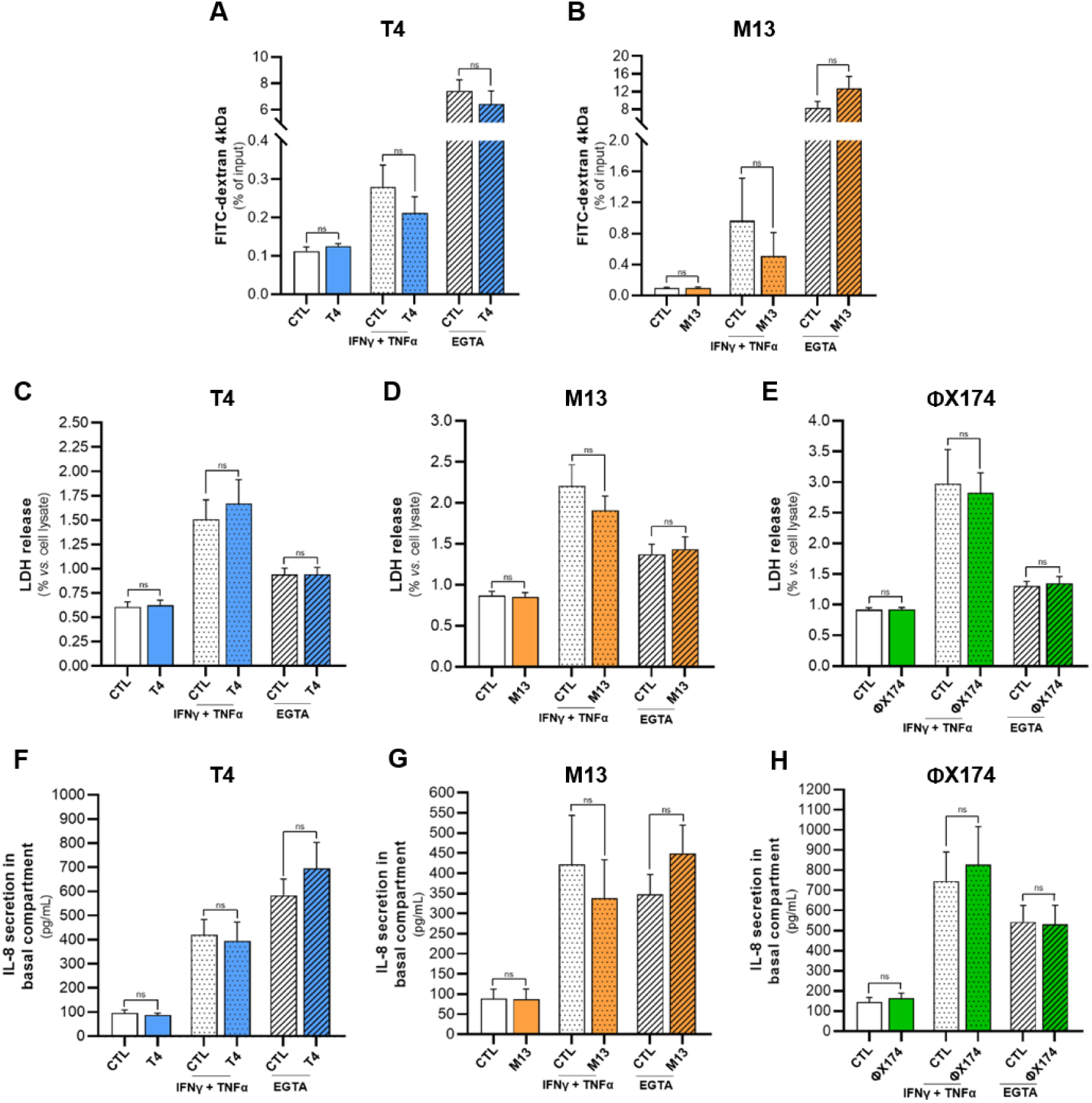
Phages do not induce hyperpermeability, cytotoxicity, or inflammation in Caco-2/TC7 cells under compromised barrier conditions. **(A-B)** FITC-Dextran 4 kDa (FD4) passage from the apical to the basal compartment during 24 h in presence of **(A)** T4, or **(B) M13** phages relative to control (CTL) condition, under healthy, and compromised barrier conditions (IFNγ + TNFα, or EGTA). **(C-E)** Release of lactate dehydrogenase (LDH) from cells after 24 h in presence of **(C)** T4, **(D)** M13, or **(E)** ΦX174 phages relative to control (CTL) condition, under healthy, and compromised barrier conditions (IFNγ + TNFα, or EGTA). (F-H) Secretion of pro-inflammatory cytokine IL-8 by the cells into the basal compartment after 24 h in presence of **(F)** T4, **(G)** M13, or **(H)** ΦX174 phages relative to control (CTL) condition, under healthy, and compromised barrier conditions (IFNγ + TNFα, or EGTA). For all these experiments, a minimum of three independent experiments [n ≥ 3] were performed.

**Figure S9:**
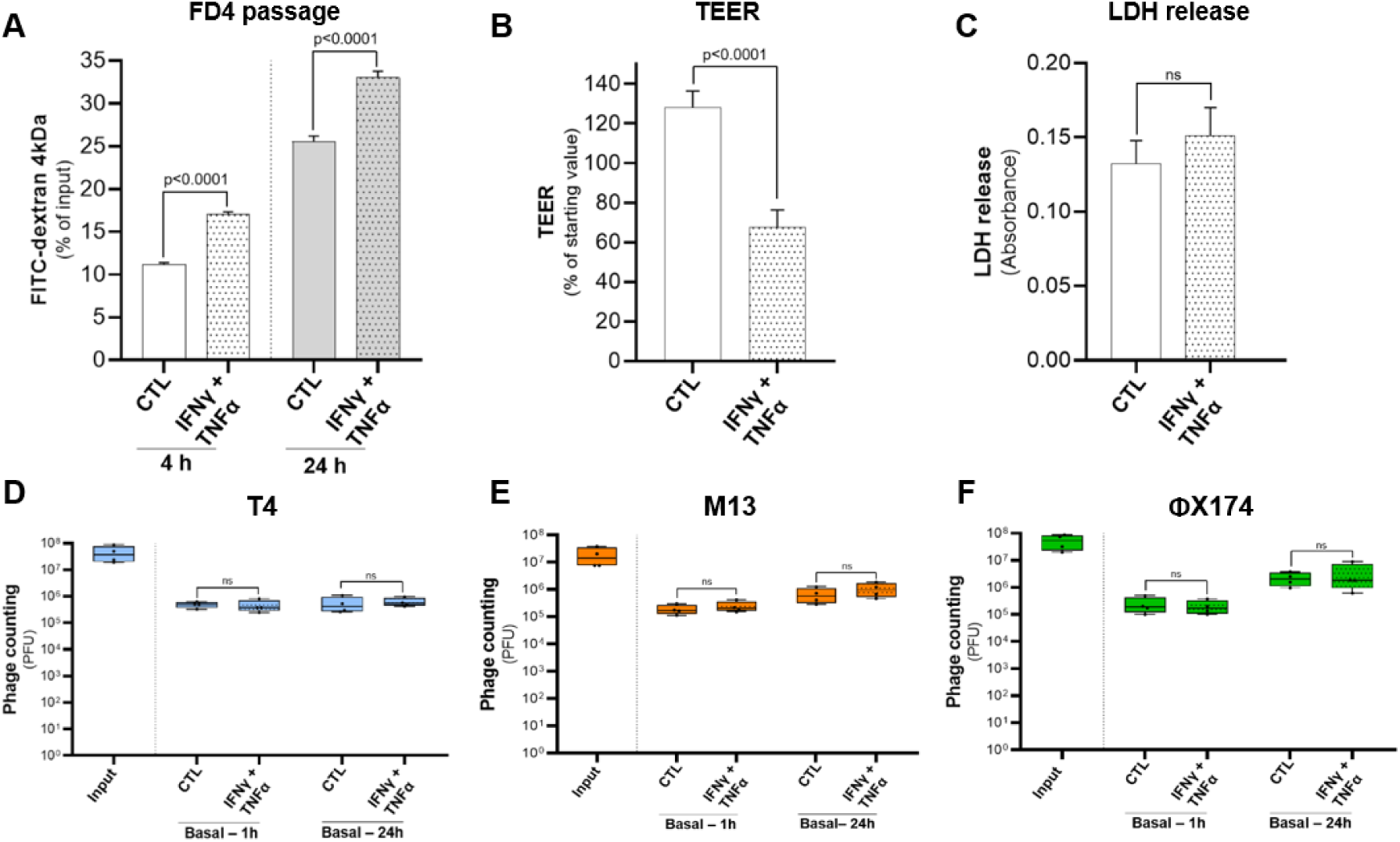
Phage translocation is not impacted by inflammatory condition in HUVECs cells. **(A)** FITC-Dextran 4 kDa (FD4) passage from the apical to the basal compartment after 22 h (4 h of FD4 flux), and 42 h (4 h of FD4 flux) of IFNγ + TNFα treatment. (B) Measure of transendothelial electrical resistance (TEER) after 18 h of IFNγ + TNFα treatment. Values are expressed as a percentage relative to the starting value for each well and then normalized to the control condition. **(C)** Release of lactate dehydrogenase (LDH) from cells after 42 h of IFNγ + TNFα treatment. **(D-F)** Titration of **(D)** T4, **(E)** M13, and **(F)** ΦX174 phages in the basal compartment (basal) after 24 h of incubation. The input represents the phages added in the apical compartment at the beginning of the experiment. Conditions include healthy barrier condition (CTL) relative to compromised barriers conditions induced by IFNγ and TNFα. Each dot in the figure represents data from one well. For (A), (B), (C)_T_ experiments were pooled from control groups used in independent experiments involving T4, M13, and ΦX174 phages [n = 9], For (D), (E), (F), three independent experiments [n = 3] were performed. Data are represented as mean ± SEM.

**Figure S10:**
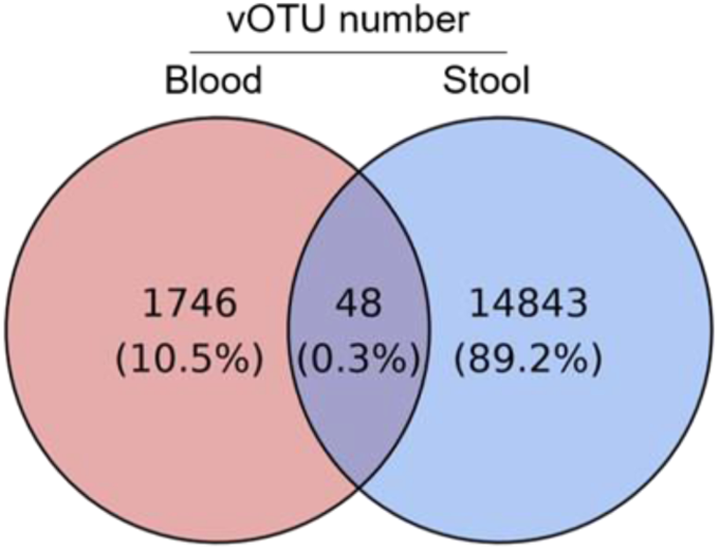
Venn diagram of the vOTUs identified in blood and stool samples from all participants. The total dataset represent 16,637 vOTUs detected tn the 29 participants of the cohort.

**Figure S11:**
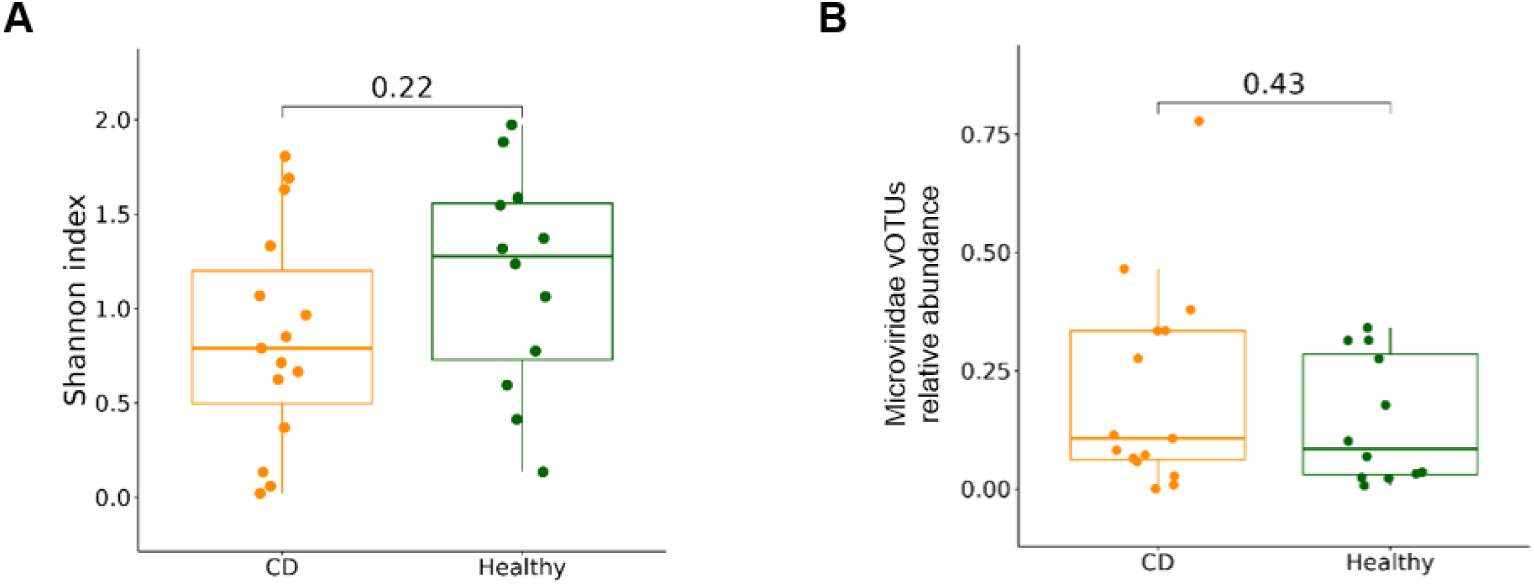
Diversity, and relative abundance of fecal *Microviridae* in CD patients, and healthy subjects. **(A)** Alpha diversity of *Microviridae* in stool samples, quantified using Shannon index, and categorized by disease status. **(B)** Overall relative abundance of *Microviridae* vOTUs in stool samples, separated by disease status.

## SUPPLEMENTARY TABLES

**Table S1:**
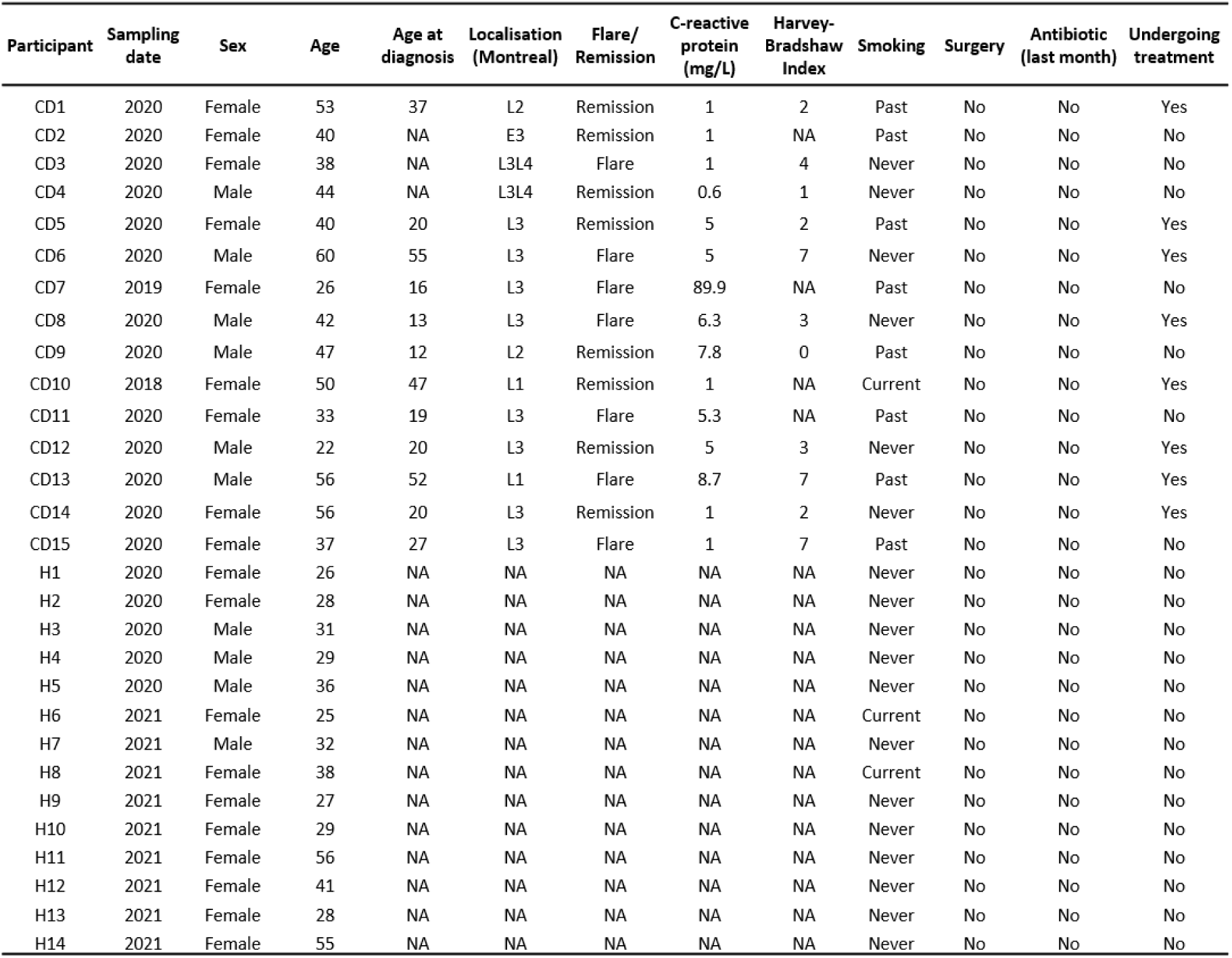
Detailed information about the participants included in the study.

| Participant | Sampling date | Sex | Age | Age at diagnosis | Localisation (Montreal) | Flare/ Remission | C-reactive protein (mg/L) | Harvey-Bradshaw Index | Smoking | Surgery | Antibiotic (last month) | Undergoing treatment |
| --- | --- | --- | --- | --- | --- | --- | --- | --- | --- | --- | --- | --- |
| CD1 | 2020 | Female | 53 | 37 | L2 | Remission | 1 | 2 | Past | No | No | Yes |
| CD2 | 2020 | Female | 40 | NA | E3 | Remission | 1 | NA | Past | No | No | No |
| CD3 | 2020 | Female | 38 | NA | L3L4 | Flare | 1 | 4 | Never | No | No | No |
| CD4 | 2020 | Male | 44 | NA | L3L4 | Remission | 0.6 | 1 | Never | No | No | No |
| CD5 | 2020 | Female | 40 | 20 | L3 | Remission | 5 | 2 | Past | No | No | Yes |
| CD6 | 2020 | Male | 60 | 55 | L3 | Flare | 5 | 7 | Never | No | No | Yes |
| CD7 | 2019 | Female | 26 | 16 | L3 | Flare | 89.9 | NA | Past | No | No | No |
| CD8 | 2020 | Male | 42 | 13 | L3 | Flare | 6.3 | 3 | Never | No | No | Yes |
| CD9 | 2020 | Male | 47 | 12 | L2 | Remission | 7.8 | 0 | Past | No | No | No |
| CD10 | 2018 | Female | 50 | 47 | L1 | Remission | 1 | NA | Current | No | No | Yes |
| CD11 | 2020 | Female | 33 | 19 | L3 | Flare | 5.3 | NA | Past | No | No | No |
| CD12 | 2020 | Male | 22 | 20 | L3 | Remission | 5 | 3 | Never | No | No | Yes |
| CD13 | 2020 | Male | 56 | 52 | L1 | Flare | 8.7 | 7 | Past | No | No | Yes |
| CD14 | 2020 | Female | 56 | 20 | L3 | Remission | 1 | 2 | Never | No | No | Yes |
| CD15 | 2020 | Female | 37 | 27 | L3 | Flare | 1 | 7 | Past | No | No | No |
| H1 | 2020 | Female | 26 | NA | NA | NA | NA | NA | Never | No | No | No |
| H2 | 2020 | Female | 28 | NA | NA | NA | NA | NA | Never | No | No | No |
| H3 | 2020 | Male | 31 | NA | NA | NA | NA | NA | Never | No | No | No |
| H4 | 2020 | Male | 29 | NA | NA | NA | NA | NA | Never | No | No | No |
| H5 | 2020 | Male | 36 | NA | NA | NA | NA | NA | Never | No | No | No |
| H6 | 2021 | Female | 25 | NA | NA | NA | NA | NA | Current | No | No | No |
| H7 | 2021 | Male | 32 | NA | NA | NA | NA | NA | Never | No | No | No |
| H8 | 2021 | Female | 38 | NA | NA | NA | NA | NA | Current | No | No | No |
| H9 | 2021 | Female | 27 | NA | NA | NA | NA | NA | Never | No | No | No |
| H10 | 2021 | Female | 29 | NA | NA | NA | NA | NA | Never | No | No | No |
| H11 | 2021 | Female | 56 | NA | NA | NA | NA | NA | Never | No | No | No |
| H12 | 2021 | Female | 41 | NA | NA | NA | NA | NA | Never | No | No | No |
| H13 | 2021 | Female | 28 | NA | NA | NA | NA | NA | Never | No | No | No |
| H14 | 2021 | Female | 55 | NA | NA | NA | NA | NA | Never | No | No | No |

**Table S2:** Number of pairs of reads per sample. The number of pairs obtained after sequencing is indicated in the "Raw” column. The number of pairs remaining after quality-filtering, and removal of non-human reads is indicated in the "Clean and non-human" column. The samples with less than 100,000 pairs of clean, and non-human reads were excluded from the analysis.

| SampleName | Status | Sample Type | Raw | Clean and non-human |
| --- | --- | --- | --- | --- |
| H1_F | Healthy | Stool | 4 557 777 | 3 302 598 |
| H2_F | Healthy | Stool | 5 141 556 | 3 538 237 |
| H3_F | Healthy | Stool | 4 575 463 | 3 314 041 |
| H4_F | Healthy | Stool | 5 006 463 | 3 702 841 |
| H5_F | Healthy | Stool | 6 630 408 | 4 868 696 |
| H6_F | Healthy | Stool | 6 043 597 | 4 533 278 |
| H7_F | Healthy | Stool | 149 | 113 |
| H8_F | Healthy | Stool | 12 607 586 | 10 702 899 |
| H9_F | Healthy | Stool | 4 957 375 | 4 008 417 |
| H10_F | Healthy | Stool | 161 | 122 |
| H11_F | Healthy | Stool | 47 423 047 | 36 124 946 |
| H12_F | Healthy | Stool | 3 848 503 | 3 021 191 |
| H13_F | Healthy | Stool | 3 944 094 | 3 101 396 |
| H14_F | Healthy | Stool | 10 220 371 | 8 099 646 |
| CD1_F | Crohn | Stool | 3 398 455 | 2 610 231 |
| CD2_F | Crohn | Stool | 11 438 302 | 8 205 616 |
| CD3_F | Crohn | Stool | 27 092 512 | 19 524 928 |
| CD4_F | Crohn | Stool | 18 270 751 | 12 128 024 |
| CD5_F | Crohn | Stool | 8 453 588 | 5 729 131 |
| CD6_F | Crohn | Stool | 4 522 582 | 3 789 808 |
| CD7_F | Crohn | Stool | 2 985 683 | 2 478 116 |
| CD8_F | Crohn | Stool | 10 219 909 | 8 279 831 |
| CD9_F | Crohn | Stool | 7 091 001 | 5 360 552 |
| CD10_F | Crohn | Stool | 9 143 203 | 7 378 428 |
| CD11_F | Crohn | Stool | 9 290 609 | 7 084 801 |
| CD12_F | Crohn | Stool | 6 350 692 | 5 222 735 |
| CD13_F | Crohn | Stool | 7 983 876 | 6 487 386 |
| CD14_F | Crohn | Stool | 636 447 | 498 993 |
| CD15_F | Crohn | Stool | 3 500 715 | 2 724 997 |
| H1_B | Healthy | Blood | 25 145 743 | 6 790 645 |
| H2_B | Healthy | Blood | 13 872 445 | 5 336 467 |
| H3_B | Healthy | Blood | 17 469 863 | 6 122 420 |
| H4_B | Healthy | Blood | 16 038 090 | 3 433 116 |
| H5_B | Healthy | Blood | 19 342 807 | 5 840 315 |
| H6_B | Healthy | Blood | 1 916 528 | 185 563 |
| H7_B | Healthy | Blood | 2 347 924 | 837 299 |
| H8_B | Healthy | Blood | 2 733 222 | 788 996 |
| H9_B | Healthy | Blood | 2 689 347 | 723 438 |
| H10_B | Healthy | Blood | 2 486 126 | 573 092 |
| H11_B | Healthy | Blood | 1 397 921 | 160 240 |
| H12_B | Healthy | Blood | 3 711 044 | 691 480 |
| H13_B | Healthy | Blood | 2 312 559 | 68 548 |
| H14_B | Healthy | Blood | 5 506 340 | 2 713 132 |
| CD1_B | Crohn | Blood | 25 941 062 | 8 696 199 |
| CD2_B | Crohn | Blood | 14 183 825 | 8 720 837 |
| CD3_B | Crohn | Blood | 29 669 303 | 14 833 794 |
| CD4_B | Crohn | Blood | 20 129 214 | 11 955 938 |
| CD5_B | Crohn | Blood | 16 967 073 | 10 236 927 |
| CD6_B | Crohn | Blood | 4 631 975 | 1 917 021 |
| CD7_B | Crohn | Blood | 2 048 220 | 12 333 |
| CD8_B | Crohn | Blood | 7 197 457 | 3 434 820 |
| CD9_B | Crohn | Blood | 6 813 381 | 3 293 988 |
| CD10_B | Crohn | Blood | 6 542 891 | 3 338 429 |
| CD11_B | Crohn | Blood | 7 374 010 | 2 778 364 |
| CD12_B | Crohn | Blood | 9 614 014 | 4 117 958 |
| CD13_B | Crohn | Blood | 5 606 138 | 2 207 758 |
| CD14_B | Crohn | Blood | 5 555 652 | 2 052 908 |
| CD15_B | Crohn | Blood | 8 436 207 | 4 380 873 |
| Microtube |  |  | 5 242 956 | 2 930 415 |
| BloodCollectionTube |  |  | 10 853 654 | 4 427 872 |

**Table S3:** Percentage of translocated phages across intestinal tissue in healthy mice. Percentage of phages that translocated after 90 min of addition in the apical compartment using the Ussing chamber system, normalized relative to the input. For each phage, three ileum or colon tissue samples were taken from two or three individual mice.

| Phage | Ileum<br>(Basal / Input)<br>Ratio in % | Colon<br>(Basal / Input)<br>Ratio in % |
| --- | --- | --- |
| **T4** | 0.0008730 (± 0.001496) | 0.0004333 (± 0.0005859) |
| **M13** | 9.348e-005 (± 0.0001615) | 1.207e-005 (± 1.987e-005) |
| **ΦX174** | 0.0004167 (± 0.0007217) | 0 |
| **p-value** | ns | ns |

**Table S4:** Phages do not induce hyperpermeability of Caco-2/TC7 cells in compromised barrier conditions. Measure of transepithelia I electrical resistance (TEER) after 24 h of incubation with medium containing no phages (CTL), or the indicated phages, under healthy, and compromised barrier conditions (IFNγ + TNFα or EGTA treatment). A minimum of three independent experiments [n ≥ 3] were performed. Values are expressed as a percentage relative to the starting value for each well, and then compared to the control condition.

| TEER (% of starting value) |  |  |  |  |  |  |
| --- | --- | --- | --- | --- | --- | --- |
|  | CTL | T4 | CTL | M13 | CTL | ΦX174 |
|  | 100<br>(± 3.643) | 106.2<br>(± 6.758) | 100<br>(± 9.452) | 101.7<br>(± 14.86) | 100<br>(± 3.649) | 114.3<br>(± 14.49) |
| *p-value* | p>0.9999 |  | p>0.9999 |  | p>0.9999 |  |
| IFNγ +<br>TNFα | 97.27<br>(± 7.581) | 105.8<br>(± 16.78) | 90.77<br>(± 10.82) | 89.99<br>(± 7.991) | 91.35<br>(± 10.91) | 92.82<br>(± 7.656) |
| *p-value* | p>0.9999 |  | p>0.9999 |  | p>0.9999 |  |
| EGTA | 48.04<br>(± 13.02) | 58.29<br>(± 20.81) | 31.06<br>(± 16.16) | 22.72<br>(± 11.42) | 45.9<br>(± 28.30) | 46.80<br>(± 30.25) |
| *p-value* | p>0.9999 |  | p>0.9999 |  | p>0.9999 |  |

